# Chromosome remodelling by SMC/Condensin in *B. subtilis* is regulated by Soj/ParA during growth and sporulation

**DOI:** 10.1101/2021.12.18.473321

**Authors:** David M. Roberts, Anna Anchimiuk, Tomas G. Kloosterman, Heath Murray, Ling Juan Wu, Stephan Gruber, Jeff Errington

## Abstract

SMC complexes, loaded at ParB-*parS* sites, are key mediators of chromosome organization in bacteria. ParA/Soj proteins interact with ParB/Spo0J in a pathway involving ATP-dependent dimerization and DNA binding, leading to chromosome segregation and SMC loading. In *Bacillus subtilis*, ParA/Soj also regulates DNA replication initiation, and along with ParB/Spo0J is involved in cell cycle changes during endospore formation. The first morphological stage in sporulation is the formation of an elongated chromosome structure called an axial filament. We now show that a major redistribution of SMC complexes drives axial filament formation, in a process regulated by ParA/Soj. Unexpectedly, this regulation is dependent on monomeric forms of ParA/Soj that cannot bind DNA or hydrolyse ATP. These results reveal a new role for ParA/Soj proteins in the regulation of SMC dynamics in bacteria, and yet further complexity in the web of interactions involving chromosome replication, segregation, and organization, controlled by ParAB and SMC.

## Introduction

The stable inheritance of chromosomes is fundamental to virtually all cells. In bacteria, the lack of an overt mitotic spindle raises intriguing questions about their mechanisms of chromosome segregation and offers the possibility of targeting the process with selectively toxic antibiotics. Endospore formation (sporulation) in the Gram-positive bacterium *Bacillus subtilis* has proved a powerful experimental system for studying chromosome segregation in bacteria. In the early stages of sporulation, cells undergo a highly asymmetric cell division that requires the chromosomes to reorganize into an axial filament, in which the chromosomes radiate along the entire length of the cell and become anchored to opposite cell poles (Figure 1A) (Glaser et al., 1997; Webb et al., 1997; Wu and Errington, 1998).

**Figure 1.**
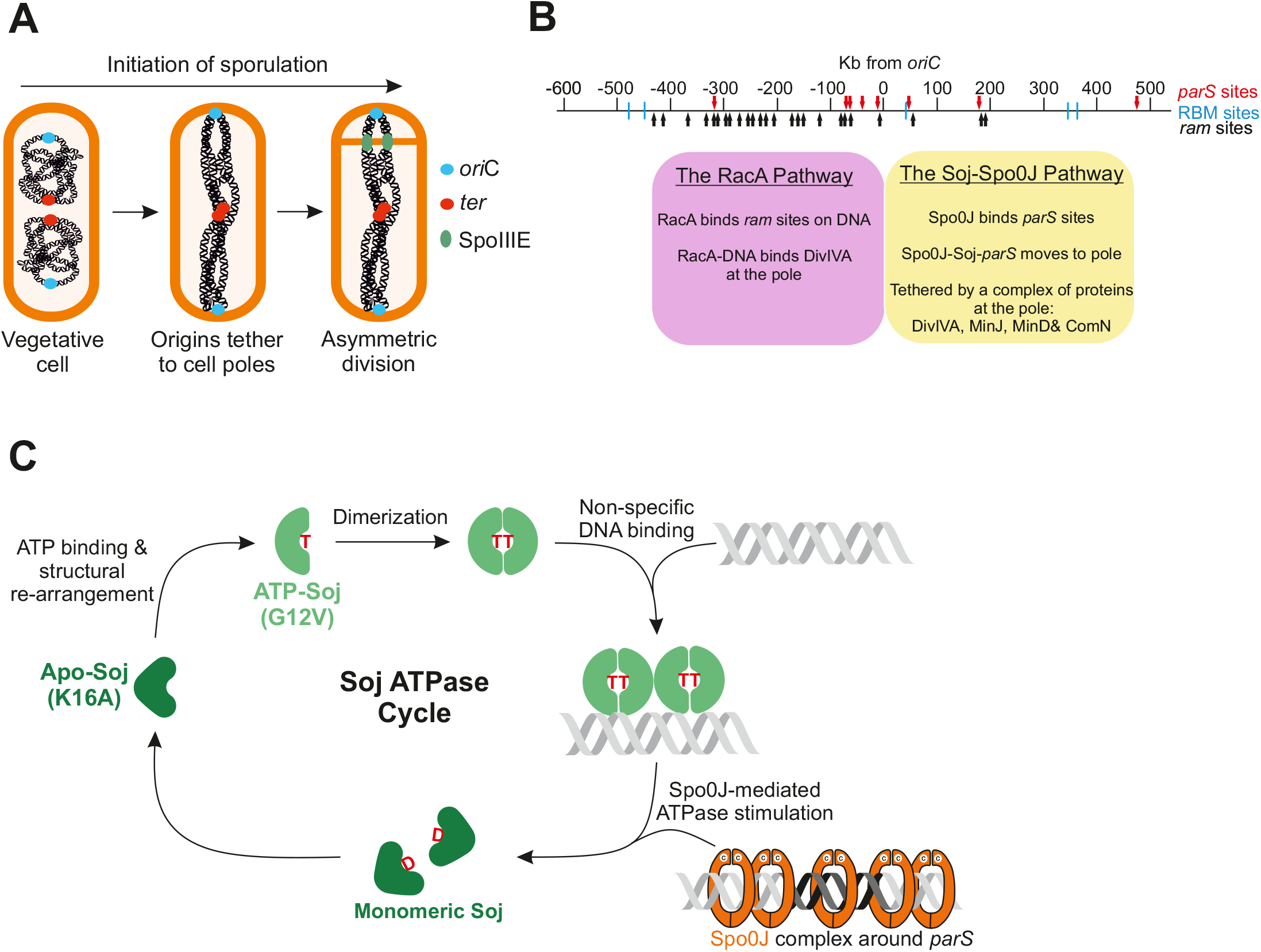
Chromosome dynamics during early sporualtion. **A)** Schematic representation of the changes to chromosome segregation during early sporulation. Chromosome origin regions (blue circles) become anchored to opposite cell poles, while the terminus region (red circles) remain associated at mid-cell. This is followed by asymmetric division that defines the small prespore and large mother cell. SpoIIIE (green circles) resides in the asymmetric septum to prevent DNA scission and pump the mother cell localized portion of the bisected chromosome into the prespore. **B)** Schematic showing the major features of the origin region required for anchoring the chromosome to the pole during sporulation. There are two redundant pathways involved in capturing the origin. The first is the sporulation specific RacA pathway, which is centred around *ram* sites (black arrows) to the left of *oriC*. The second pathway is centred around the *parS* sites at *oriC* (red arrows). Spo0J binds *parS*, and along with Soj and a complex of polar proteins, tethers the origin region to the pole. Blue lines = RBM sites.

Upon polar division, one of the chromosomes in the axial filament becomes bisected by the closing septum such that the majority of the chromosome is located in the large compartment (outside the spore) (Wu and Errington, 1994, 1998). SpoIIIE (FtsK in other organisms) binds the bisected chromosome, preventing its scission, and then translocates >3 Mbp of DNA through a pore in the septum to complete chromosome partitioning into the small prespore (Bath et al., 2000; Ben-Yehuda and Losick, 2002; Burton et al., 2007; Fiche et al., 2013; Wu and Errington, 1997; Yen Shin et al., 2015). Following polar division, the prespore and larger mother cell activate distinct gene expression programmes, with the transcription factor σ^F^ becoming specifically activated in the prespore (Harry et al., 1995; Lewis et al., 1994; Margolis et al., 1991). By placing σ^F^-specific reporters at various locations on the chromosome, a detailed map of the prespore-trapped segment was determined and found to comprise ∼1 Mbp of DNA roughly centred on the origin of replication (origin, or *oriC*) (Wu and Errington, 1994, 1998).

Similar “chromosome trapping assays” with σ^F^-responsive reporter genes were also used to identify and characterize genes required for *oriC* movement to the cell pole (Figure 1B). The sporulation specific RacA protein binds specific recognition (*ram*) sites in the chromosome clustered to the left of *oriC* (Figure 1B) (Ben-Yehuda et al., 2005; Ben-Yehuda et al., 2003; Wu and Errington, 2002, 2003). As well as binding to *ram* sites, RacA interacts with DivIVA, a landmark protein that associates with cell division sites and poles (Lenarcic et al., 2009), such that origins are physically tethered to the cell poles in a “Velcro”-like manner. The function of RacA is partially redundant to the more complex Soj-Spo0J system (Kloosterman et al., 2016; Sharpe and Errington, 1996; Wu and Errington, 2003). *soj* and *spo0J* are the *B. subtilis* homologues of the *parA* and *parB* genes present in most bacteria and on many low copy number plasmids, and which are thought to play a direct role in chromosome and plasmid segregation (Baxter and Funnell, 2014; Funnell, 2016; Jalal and Le, 2020). The Soj-Spo0J system acts at *parS* sites – specific loading sites for Spo0J that are mainly located close to *oriC*, the single origin of bacterial chromosome replication (Figure 1B) (Lin and Grossman, 1998; Livny et al., 2007; Murray et al., 2006). Genetic experiments have revealed several other genes required for axial filament formation including, in addition to *divIVA, comN, minJ, minD*, and *sirA* (Autret and Errington, 2003; Duan et al., 2016; Kloosterman et al., 2016). Combined elimination of the RacA and Soj-Spo0J pathways virtually abolishes prespore chromosome trapping, suggesting that they have complementary roles in mediating polar origin movement (Kloosterman et al., 2016; Wu and Errington, 2003). A third system has been implicated in coordinating polar chromosome segregation and the subsequent asymmetric cell division. In this system, RefZ simultaneously interacts with RefZ-binding motifs (RBMs) on the DNA and the major cell division protein, FtsZ, although the precise function of this system remains unclear (Brown et al., 2019; Miller et al., 2015; Wagner-Herman et al., 2012).

ParB/Spo0J proteins are DNA binding CTP hydrolases (Jalal et al., 2020; Osorio-Valeriano et al., 2019; Soh et al., 2019). ParB-CTP dimers have an open configuration, which upon interaction with *parS* sequences, specifically clamp shut around the DNA. Closed ParB-dimers are then released from *parS*, allowing local spreading via lateral (and possibly bridging) interactions, before CTP hydrolysis and release from DNA (Graham et al., 2014; Jalal et al., 2020; Murray et al., 2006; Osorio-Valeriano et al., 2021; Osorio-Valeriano et al., 2019; Soh et al., 2019; Taylor et al., 2015; Taylor et al., 2021). ParA/Soj proteins are Walker ATPases. Binding of ATP to Apo-ParA drives a structural change that allows dimerization, and ParA-dimers can bind DNA non-specifically (Figure 1C) (Hester and Lutkenhaus, 2007; Leonard et al., 2005; McLeod and Spiegelman, 2005), although the extent of ParA-ATP dimer binding on the genome varies amongst bacterial species. For example, in *Caulobacter crescentus* ParA dimers bind non-specifically over the entire chromosome (Lim et al., 2014; Ptacin et al., 2010; Toro et al., 2008). By contrast, ParA/Soj binding is largely restricted to the *oriC* region in *B. subtilis* (Murray and Errington, 2008), unless ParB/Spo0J is absent, in which case ParA/Soj dimers accumulate and bind more widely over the chromosome (Marston and Errington, 1999; Murray and Errington, 2008; Quisel et al., 1999). The extreme N-terminal region of ParB/Spo0J triggers ATP hydrolysis in ParA/Soj dimers, resulting in dissociation of the dimer and release from DNA (Gruber and Errington, 2009; Scholefield et al., 2011; Surtees and Funnell, 1999).

One proposed mechanism for ParA/ParB mediated chromosome segregation is the DNA relay system of *C. crescentus*, which drives directed movement of one chromosome origin from the stalked to the flagellated pole (Lim et al., 2014). In this system, ParA-ATP dimers bind non-specifically across the chromosome. Newly replicated ParB-*parS* complexes then interact with nearby ParA dimers on the DNA, driving ParA-ATP hydrolysis and release from the chromosome, followed by ParB-*parS* interaction with another ParA dimer. By sequentially repeating this process, the ParB-*parS* complex follows the retreating “cloud” of ParA on the DNA. A similar mechanism has been proposed for the segregation of ParABS plasmids (Hu et al., 2015; Hwang et al., 2013; Lim et al., 2014; Surovtsev et al., 2016a; Surovtsev et al., 2016b; Taylor et al., 2021; Vecchiarelli et al., 2013; Vecchiarelli et al., 2014).

In *B. subtilis, soj* and *spo0J* deletions only elicit mild effects on chromosome segregation in vegetative cells. However, Soj and Spo0J clearly have important roles in a variety of crucial cell cycle processes. We previously showed that monomer and dimer forms of Soj have opposing effects on the regulation of DNA replication (inhibiting or promoting it, respectively) through direct interactions with the master initiator of DNA replication, DnaA (Murray and Errington, 2008; Scholefield et al., 2012; Scholefield et al., 2011). It then emerged that Spo0J contributes to chromosome segregation by recruiting and loading the bacterial SMC/Condensin complex (Gruber and Errington, 2009; Sullivan et al., 2009). SMC complexes align and juxtapose left and right chromosome arms, and prevent chromosome tangling, as they travel from their Spo0J-*parS* loading site to the terminus region, after which they are specifically unloaded by XerD (Gruber et al., 2014; Karaboja et al., 2021; Wang et al., 2017; Wang et al., 2018; Wang et al., 2015; Wang et al., 2014). *spo0J* mutants are also deficient in sporulation because in the absence of Spo0J, Soj accumulates in the ATP-dimer form that promotes DNA over-replication (Murray and Errington, 2008). This blocks sporulation via a checkpoint mechanism involving an inhibitor of sporulation called *sda* (Burkholder et al., 2001; Veening et al., 2009). Thus, Soj and Spo0J have a pivotal role in multiple cell cycle events coupling DNA replication initiation, origin separation, chromosome segregation and sporulation. Moreover, although SMC complexes are conserved from bacteria to man, and have been extensively studied in a variety of systems, Spo0J was the first sequence-specific loading factor to be identified (Gruber and Errington, 2009; Sullivan et al., 2009).

The structural change in chromosome configuration associated with the early stages of sporulation was first recognized more than fifty years ago and named the axial filament (Ryter, 1965). Discovery that *oriC* regions of the chromosome are specifically trapped in the prespore compartment, and of factors required for their positioning, have led to the assumption that *oriC* migration is the main driving force in axial filament formation. Here, we report a series of surprising findings that change our understanding of axial filament formation and the roles of ParAB and SMC complexes. First, we show that in *B. subtilis*, ParA/Soj is active in chromosome segregation in its <u>monomeric</u>, non-DNA binding form. Furthermore, we show that ParA/Soj monomers can exist in two functional states, likely corresponding to the Apo- and ATP-bound forms, and that the major functional difference between these forms lies in the regulation of SMC complex redistribution at ParB/Spo0J-*parS* sites. Finally, we show that axial filament formation is accompanied by a major redistribution in SMC complexes on the chromosome, and that the monomer forms of ParA/Soj control this positioning, thus providing new mechanistic insights into chromosome dynamics in bacteria.

## Results

### Wild type Soj protein appears mainly monomeric

To gain insights into the function of Soj in chromosome segregation during sporulation we prepared a new fusion of Soj to the mNeonGreen (mNG) fluorescent protein (Shaner et al., 2013), which is unrelated to GFP, and is brighter than previously used GFP-Soj fusions. As shown in Figure 2 (top row, left panel) early sporulating cells expressing mNG-Soj as the only functional copy of Soj showed a complex pattern with at least three detectable elements: a diffuse background signal throughout the cell; a septal signal, which appeared to be associated with recently completed vegetative septa; and an extreme polar focus, reminiscent of the localization of proteins DivIVA, MinD and ComN, involved in the ORI-zone trapping system, or “polar complex”, described previously (Kloosterman et al., 2016).

**Figure 2.**
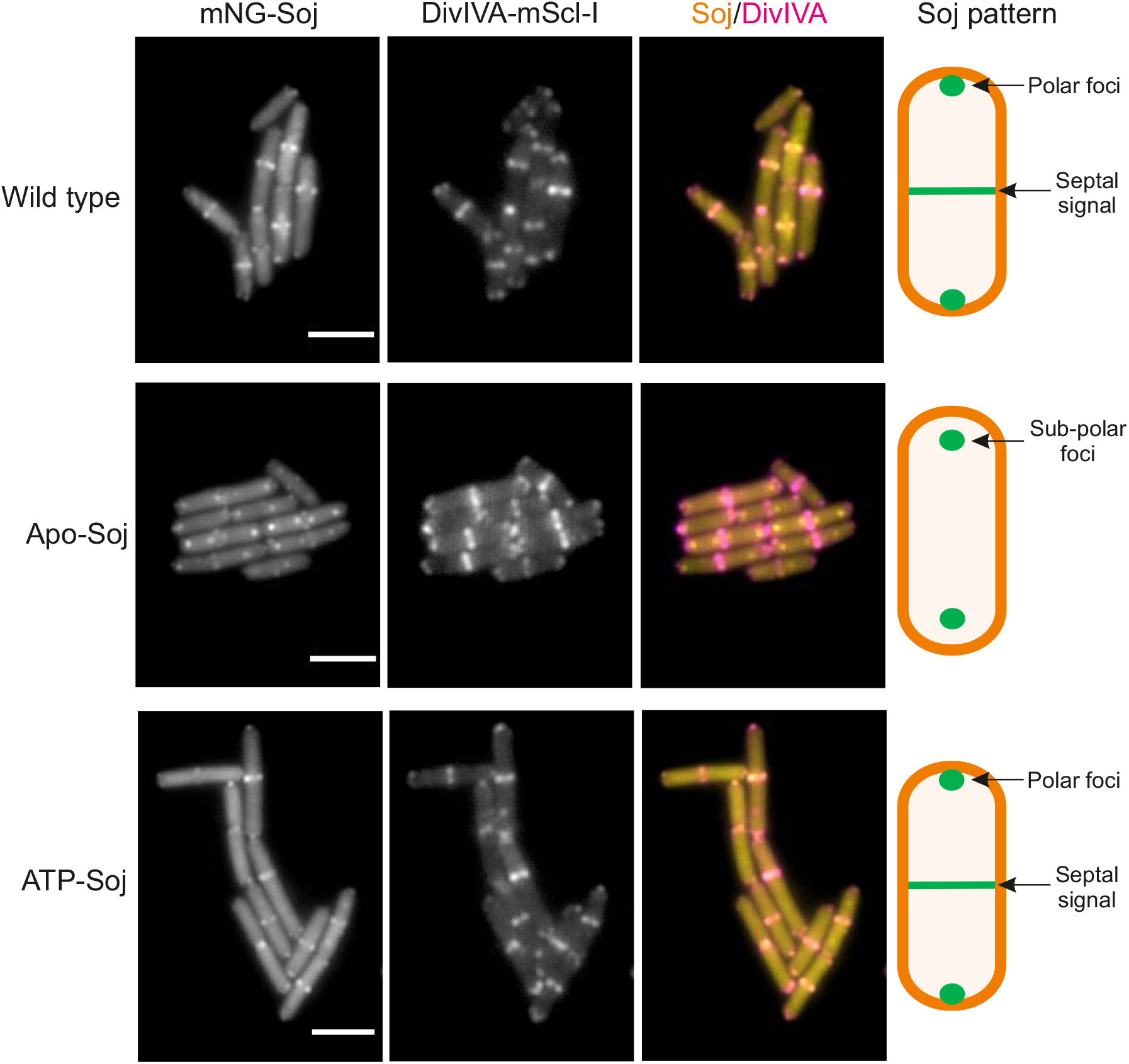
Co-localization of Soj and DivIVA. Representative images showing the localization patterns of the mNG-Soj mutants and DivIVA-mScl-I (Apo-Soj = *soj(K16A)*; ATP-Soj = *soj(G12V)*). Imagges were acquired 80 mins after re-suspension in sporulation salts. Scale bar = 3 μm. The schematic beside each set of images shows the Soj localization pattern.

Previous work has shown that amino acid substitutions K16A and G12V trap Soj in monomeric forms that are either empty or bound to ATP, respectively (Figure 1C) (Quisel et al., 1999; Scholefield et al., 2011). Hereafter we refer to Soj(K16A) as Apo-Soj and Soj(G12V) as ATP-Soj. As shown in Figure 2 (bottom row, left panel), localization of ATP-Soj was essentially indistinguishable from the wild type (WT) fusion protein, with prominent extreme polar spots and septal bands. Apo-Soj also exhibited foci near cell poles but they appeared less strictly polar (sub-polar) than the WT or ATP-Soj forms (see below), and strikingly mid-cell bands were mainly missing (Figure 2, middle row, left panel). This was consistent with the notion that Apo-Soj does not interact with the putative sporulation “polar complex”. All three patterns were quite unlike the nucleoid-like patterns obtained with WT Soj in the absence of Spo0J, when the protein accumulates in its dimeric form, which binds DNA non-specifically (Supplementary Figure 1A).

To test more directly for possible interactions with the polar complex (Kloosterman et al., 2016), the strains containing the mNG-Soj fusion proteins also contained DivIVA-mScarlet-I (DivIVA-mScl-I). DivIVA is a major hub protein at the cell pole, and the likely upstream target for the polar complex (Kloosterman et al., 2016). Figure 2 shows representative images that suggest that WT and ATP-Soj septal bands (representing new cell poles) and extreme polar foci (older cell poles) largely co-localized with DivIVA-mScl-I. Strikingly, the Apo-Soj spots described as sub-polar above indeed failed to overlay with the DivIVA-mScl-I signal. To interrogate this further, we determined the relative cellular position of Soj and DivIVA foci in every cell, and then plotted these positions as fluorescence profiles in which each cell is stacked left to right by increasing cell length (Supplementary Figure 1B-D). This confirmed that mNG fusions of WT and ATP-Soj co-localised tightly with DivIVA-mScl-I at mid-cell septa, as determined by the tight overlay of foci generating white spots in the merged profile (Supplementary Figure 1B and D). Apo-Soj clearly showed very little overlay with the DivIVA signal at central septa. In general, the Apo-Soj foci appeared sub-polar at both outer (old) cell poles and medial (new) septa (Supplementary Figure 1C). These data were further supported by quantitative analysis of the overlap in fluorescent signals between Soj and DivIVA using co-localization analysis (Supplementary Figure 2).

Together these data support the idea that wild type Soj is mainly monomeric in sporulating *B. subtilis* (as well as vegetative cells; (Murray and Errington, 2008)), since the localisation pattern of ATP-Soj mirrored that of WT Soj, and was distinct from that observed when *spo0J* was deleted. Furthermore, both wild type and ATP-Soj co-localised substantially with DivIVA, especially at mid cell, consistent with the notion that Soj is recruited to the polar complex. In contrast, Apo-Soj forms sub-polar foci, which generally do not co-localize with DivIVA, especially at new cell poles.

### ATP-Soj can support ORI-zone trapping during sporulation

Previous work has shown that a *soj* null mutant is deficient in the ORI-trapping function during sporulation (Kloosterman et al., 2016; Wu and Errington, 2003). Given that the wild type protein appears mainly monomeric *in vivo* (Figure 2), we wondered whether the Soj monomer mutants might retain ORI-trapping activity. We took advantage of an assay based on the expression of prespore-specific reporter genes placed in the “ARM” or “ORI” regions of the chromosome (Kloosterman et al., 2016; Sullivan et al., 2009) (Figure 3A). When cells bearing a “frozen” mutant of the septum localised SpoIIIE translocase are induced to sporulate they form an asymmetric septum as normal. However, the bulk of the chromosome, that lies outside the prespore compartment when the polar septum forms, remains in the mother cell. Segments of DNA that are correctly located in the prespore can be detected if they contain a reporter gene dependent on the prespore-specific sigma factor, σ^F^ (Sullivan et al., 2009; Wu and Errington, 1994, 1997). The assay cells carried two reporter genes expressing fluorescent proteins; one located in the region of the chromosome dependent on RacA for efficient trapping (the ARM region; −418 Kb from *oriC*), and the other in the Soj-Spo0J-*parS*-dependent ORI region (−79 Kb from *oriC*) (Figure 3A) (Kloosterman et al., 2016; Sullivan et al., 2009).

**Figure 3.**
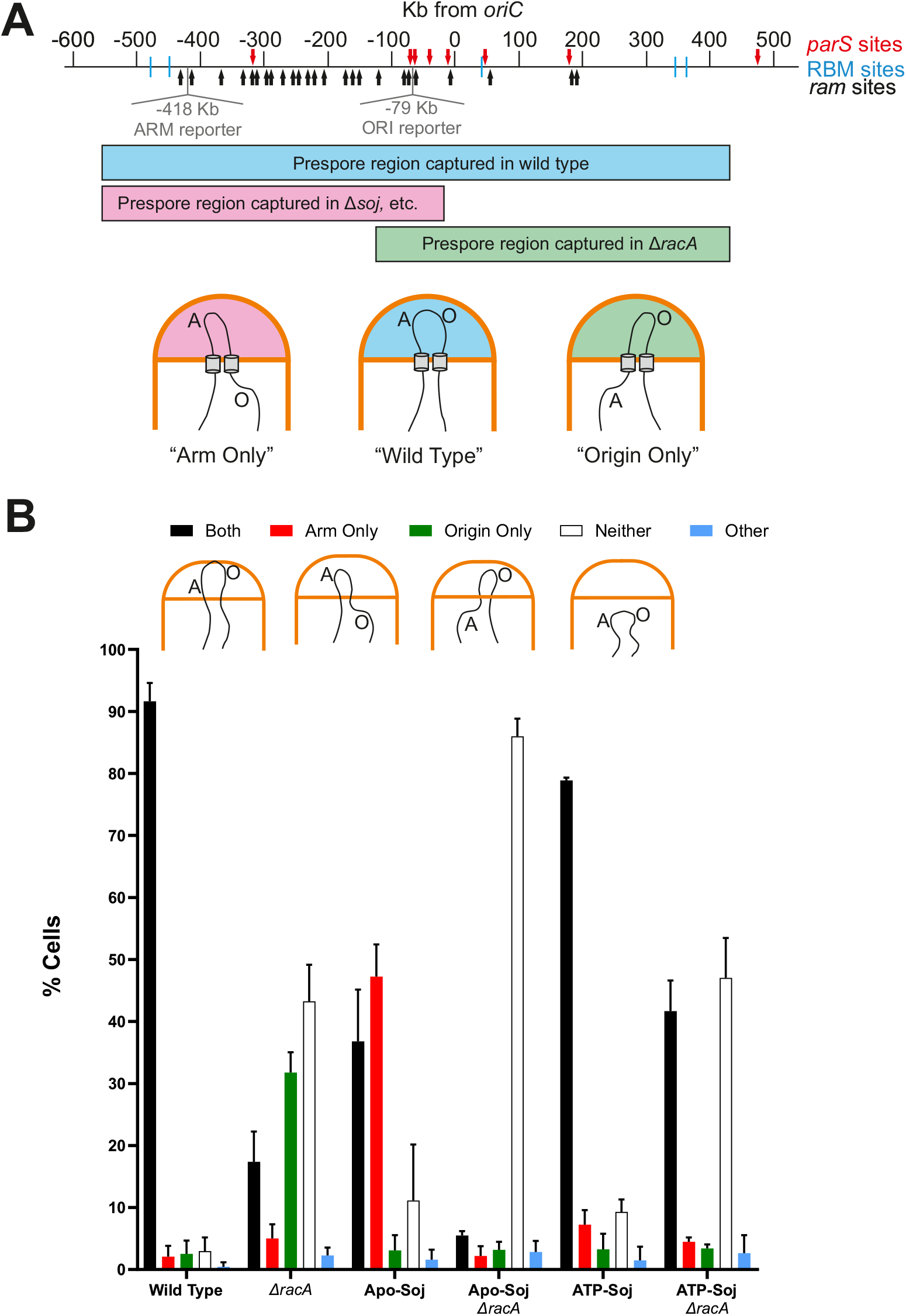
Monomeric Soj mutants have opposing effects in the chromosome trapping assay. **A)** Schematic showing the major features of the origin region required for anchoring the chromosome to the pole. There are two redundant pathways involved in origin capture. RacA binds to *ram* sites (black arrows) in the origin-proximal region, which then tethers the region to the pole. Mutations in *racA* cause an “origin only” phenotype (green). *parS* sites (red arrows) at *oriC* are the target of the Soj-Spo0J pathway that tethers the region to the pole. Mutants in *soj, spo0J* or other members of the polar complex cause an “arm only” phenotype (pink). Wild type cells successfully capture both regions (blue). The genomic position of prespore-specific reporter genes in the ORI and ARM regions are shown in grey. **B)** Chromosome trapping assay to assess the prespore localization of arm- or origin-localized fluorescent markers. At least 100 cells were scored per repeat (n = 3) and averages are shown. Error bars show standard deviations. The schematic below each category panel shows the chromosome localization pattern in that condition.

Wild type sporulating cells efficiently trapped (i.e. expressed) both ARM and ORI reporters, as expected (black bars in Figure 3B), with few cells containing only one or neither reporters. Control cells bearing a *racA* deletion showed the characteristic defect (Ben-Yehuda et al., 2003; Wu and Errington, 2003) in which less than 20% of cells correctly trapped both markers; the majority containing either only the ORI marker (“ORI-only” phenotype) or neither markers. As anticipated, cells producing Apo-Soj behaved like a *soj* null mutant (Kloosterman et al., 2016), with an excess of ARM-only capture (red bars in Figure 3B). Combining Apo-Soj with a Δ*racA* mutation resulted in an almost complete loss of trapping of either ARM or ORI markers, consistent with these mutations together abolishing both mechanisms for origin region trapping. Surprisingly, cells producing ATP-Soj were almost as proficient in trapping as wild type cells, with only a small accumulation of ARM-only or empty prespores (Figure 3B). Furthermore, when combined with Δ*racA* mutant the strain was <u>more</u> proficient in correct trapping (black bars) than the Δ*racA* single mutant (Figure 3B).

This unexpected result showed that monomeric ATP-Soj can, directly or indirectly, support movement and capturing of the origin region at the cell pole, whereas Apo-Soj cannot. Importantly, the finding that the ATP monomer form of Soj supports ORI trapping suggests that this function does not require Soj to dimerize or bind to DNA, nor does it require a functional ATPase cycle.

### Altered chromosomal distribution of SMC in monomeric mutants of *soj*

It was surprising that in the Δ*racA* background trapping of the ORI marker was more frequent with the ATP-Soj variant than for WT Soj (Figure 3B). It was conceivable that this could be due to an effect on chromosome condensation, perhaps involving altered loading of the SMC/Condensin complex at Spo0J-*parS* sites. To investigate this, we used chromatin-immunoprecipitation coupled to deep sequencing (ChIP-Seq) to plot the distribution of SMC complexes associated with the chromosome under conditions similar to the trapping assay (i.e T_3h_ sporulation and *spoIIIE36*). As shown in Supplementary Figure 3, SMC complexes were detected at sites spread around the chromosome, irrespective of the form of Soj present. As expected for wild type Soj, major peaks of SMC were associated with sites roughly corresponding to the locations of the *oriC*-proximal *parS* sites (panel A; zoomed version in panel D), which are the major loading sites for SMC complexes. However, the two mutant strains both differed from the wild type in the extent of enrichment in the *parS-oriC* region: substantially less enrichment of SMC was seen for the ATP-Soj form (Supplementary Figure 3B and E), whereas the Apo-Soj mutant had relatively more SMC over the region (Supplementary Figure 3C and F). Thus, the *soj* monomer mutations appear to affect SMC loading or dynamics, and again the Apo- and ATP-bound proteins had opposing effects.

### Apo-Soj has a dominant negative chromosome segregation defect

The above observations suggested models in which Soj regulates the loading and or release of SMC at Spo0J-*parS* sites, and that correct loading is required for ORI trapping during sporulation. If one or both proteins were trapped in forms that positively or negatively regulate SMC loading, the mutant alleles might be dominant to wild type *soj*. To test this, we engineered strains in which we could express the mutant and wild type proteins in parallel. The strain background used also carried a *dnaA(V323D)* mutation to avoid the inhibitory effects of monomeric Soj proteins (especially ATP-Soj) on the initiation of DNA replication (Murray and Errington, 2008; Scholefield et al., 2012), as well as Δ*racA* to eliminate the alternative polar segregation pathway of sporulation. These experiments revealed, unexpectedly, a clear dominant negative phenotype for Apo-Soj but not for ATP-Soj. During vegetative growth cells expressing Apo-Soj (i.e. +xylose) had a readily detectable alteration in nucleoid appearance, compared to cells expressing ATP-Soj or to either of the uninduced strains (i.e. –xylose) (Figure 4A). In particular, induction of Apo-Soj substantially reduced the number of nucleoids per cell (Figure 4B). These measurements could be influenced by changes in cell length, but the average cell length did not vary appreciably between any of the samples (Figure 4B). Thus, cells expressing Apo-Soj were delayed for nucleoid separation relative to uninduced cells or cells expressing ATP-Soj.

**Figure 4.**
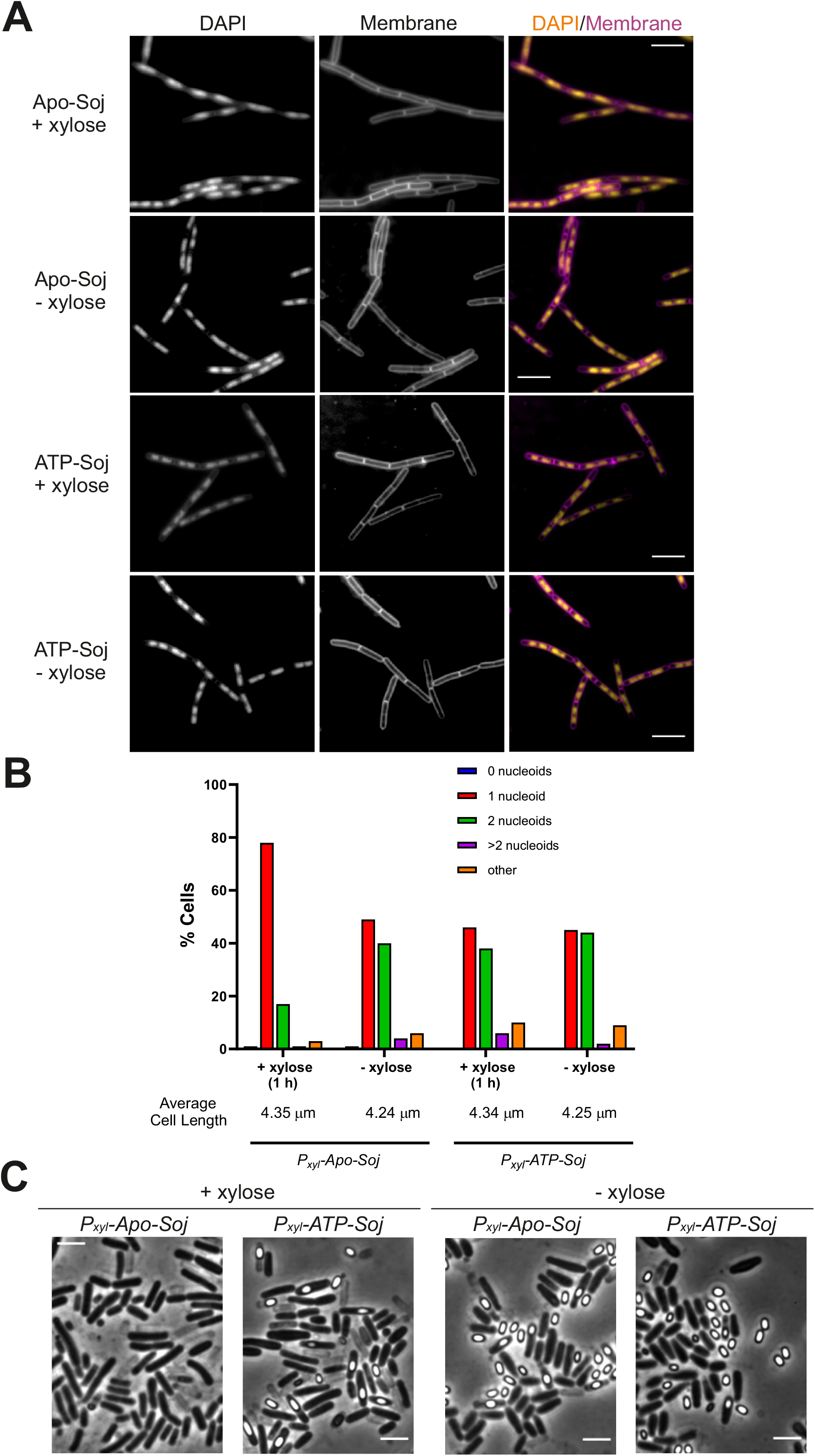
The Apo-form of Soj appears dominant negative over wild type. **A)** Representative images of vegetative cells harbouring an inducible copy of either Apo-Soj or ATP-Soj. Images were acquired 60 mins after induction of the monomer mutants using 0.5 % xylose. Scale bar = 5 μm. **B)** Bar chart showing the number of completely segregated nucleoids in the Apo- and ATP-Soj overexpression strains during vegetative growth. At least 150 cells were counted for each condition. **C)** Representative images of sporulating cells containing the inducible Apo- or ATP-Soj variant. Overexpression was induced by the addition of 0.5 % xylose (+ xylose). -xylose = no induction of the extra copy of Apo- or ATP-Soj.

When the same strains were induced to sporulate a much stronger phenotype was observed for Apo-Soj, in that induction with xylose virtually abolished sporulation (Figure 4C). Again, no significant phenotypic effect was observed for the ATP-Soj expressing strain. Detailed microscopic analysis of cells expressing Apo-Soj induced to sporulate revealed that they were blocked very early in the process, with a severe reduction in the formation of sporulation (polar) septa (as compared to uninduced Apo-Soj, or ATP-Soj in any condition) (Supplementary Figure 4A). It is well known that perturbation of chromosome replication, e.g. by the action of dimeric Soj-based stimulation of DnaA activity (Murray and Errington, 2008; Scholefield et al., 2011) can block the initiation of sporulation by a “checkpoint” mechanism involving proteins Sda, KinA and Spo0A (Burkholder et al., 2001; Veening et al., 2009). Unexpectedly, however, deletion of the checkpoint gene *sda* did not suppress the sporulation defect (Supplementary Figure 4B). We also asked whether the early sporulation-specific promoter of the *spoIIA* operon, which is dependent on activated Spo0A at the end of the above pathway (Veening et al., 2005), was expressed on induction of Apo-Soj and indeed it was (Supplementary Figure 4C). To our knowledge, this phenotype in which the early sporulation operon *spoIIA* is expressed but asymmetric cell division is blocked has only previously been reported for mutations or antibiotics inhibiting cell division directly (Beall and Lutkenhaus, 1991; Daniel et al., 1998; Stokes et al., 2005). It seems unlikely that the Apo-Soj protein would act directly on cell division. Given the effects of Apo-Soj expression on nucleoid separation described above, it seemed possible that the block in polar septation was due to incorrect positioning of the nucleoid – that is, some kind of “nucleoid occlusion” effect on cell division (Adams et al., 2015).

### Axial filament formation is associated with altered SMC distribution

The results described above suggested a model in which Soj protein regulates the loading and or sliding of SMC complexes, which then influences nucleoid configuration, i.e., formation of the axial filament, during sporulation. Correct formation of the axial filament could not only drive the origin regions towards opposite cell poles, but also configure the nucleoid to either avoid occluding the polar septum or even contribute positively to guiding septal positioning.

We took wild type cells bearing an SMC-mNG fusion and induced them to sporulate by the resuspension method. Samples taken immediately prior to resuspension (t_0_) were growing in a relatively rich medium and would be expected to have 2 to 4 copies of *oriC* (Hauser and Errington, 1995; Sharpe et al., 1998). Since SMC complexes are enriched near *oriC* due to loading at *parS* sites, 2 to 4 well-spaced foci of SMC-mNG would also be expected, and were indeed observed, in these samples (Figure 5A, E). Interestingly, however, after 1.5 h (t_90_) in sporulation medium the numbers of SMC foci increased, and in those cells with multiple foci they were distributed across the full length of the cell (Figure 5B & 5E). These time points correspond to the period when cells are progressing through the early stages of sporulation. During this process, ongoing rounds of replication are completed, generating cells with two complete chromosomes (Hauser and Errington, 1995). Meanwhile, the axial filament structure forms, in which the two *oriC* regions move to the extreme poles of the cell, while the termini remain together close to mid-cell.

**Figure 5.**
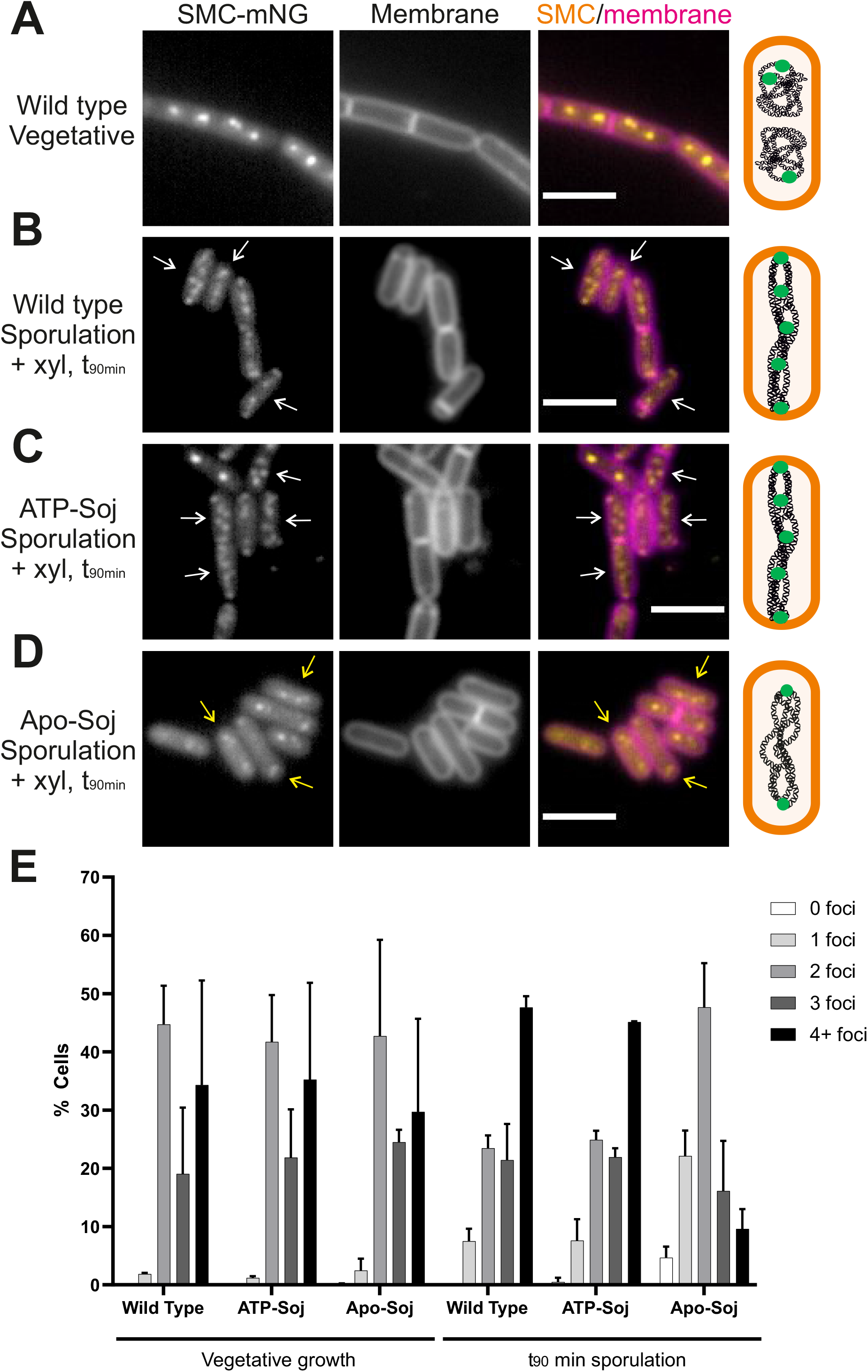
SMC distribution during sporulation. Representative images of SMC-mNG in **A)** vegetatively growing cells with wild type *soj*; **B)** cells containing wild type *soj* 90 mins after re-suspension in sporulation salts; **C)** cells containing ATP-Soj 90 mins after re-suspension in sporulation salts; **D)** cells containing Apo-Soj 90 mins after re-suspension in sporulation salts. In all cases, 0.5 % xylose was added to cells 15 mins prior to (and during) re-suspension in sporulation salts. White arrows = cells with redistributed SMC along axial filaments; yellow arrows = cells with sub-polar pairs of SMC foci. The schematic beside each set of images represents the distribution of SMC (green circles) along the chromosome in each strain. Scale bar = 3 μm. **E)** Bar chart showing the number of SMC foci detected per cell in each mutant and time point. Between 100-250 cells were counted per repeat (n = 2), with mean scores shown. Error bars show standard deviations.

These observations are consistent with previous work showing that ScpB-YFP localises along the chromosome arms during early sporulation (Wang et al., 2015). They also raised the exciting possibility that redistribution of SMC along the chromosome contributes to axial filament formation, and that Soj might play a role in regulating this process.

### Apo-Soj interferes with the redistribution of SMC protein during the early stages of sporulation

In parallel with the above experiment, we also examined the effects of ectopic expression of Apo- and ATP-Soj on SMC localization (Figure 5). Expression of ATP-Soj had little effect on the results and both SMC focus numbers and distribution were similar to those of the wild type (Figure 5C and 5E). However, xylose induction of Apo-Soj gave a strikingly different result: at t_90_ most cells still had only two widely spaced foci, presumably corresponding to the two copies of *oriC* (Figure 5D-E), suggesting that Apo-Soj interferes with the redistribution of SMC complexes needed for axial filament formation.

Although cells with the Apo-Soj mutation at the native locus (i.e. not overexpressed and no wild type protein) are not blocked for septum formation, we wondered whether they might nevertheless show a perturbation in localization of SMC. As shown in Supplementary Figure 5, these Apo-Soj mutant cells also showed a reduction in numbers of SMC foci during the early stages of sporulation, whereas cells with the ATP-Soj mutation resembled the wild type control. These findings suggested that the Apo-Soj and ATP-Soj mutant proteins have contrasting effects on the loading and/or release of SMC complexes from the origin region during sporulation.

### SMC complexes are differentially enriched at *parS* and along the chromosome arms by ATP-Soj and Apo-Soj

As well as monitoring the effects of ectopic expression of Apo- and ATP-Soj on the cellular localisation of SMC by microscopy during axial filament formation (i.e. t_90_) (Figure 5), we also conducted whole genome ChIP-Seq at these time points to establish the distribution of SMC across the entire genome in these cells (Figure 6 and Supplementary Figure 6).

**Figure 6.**
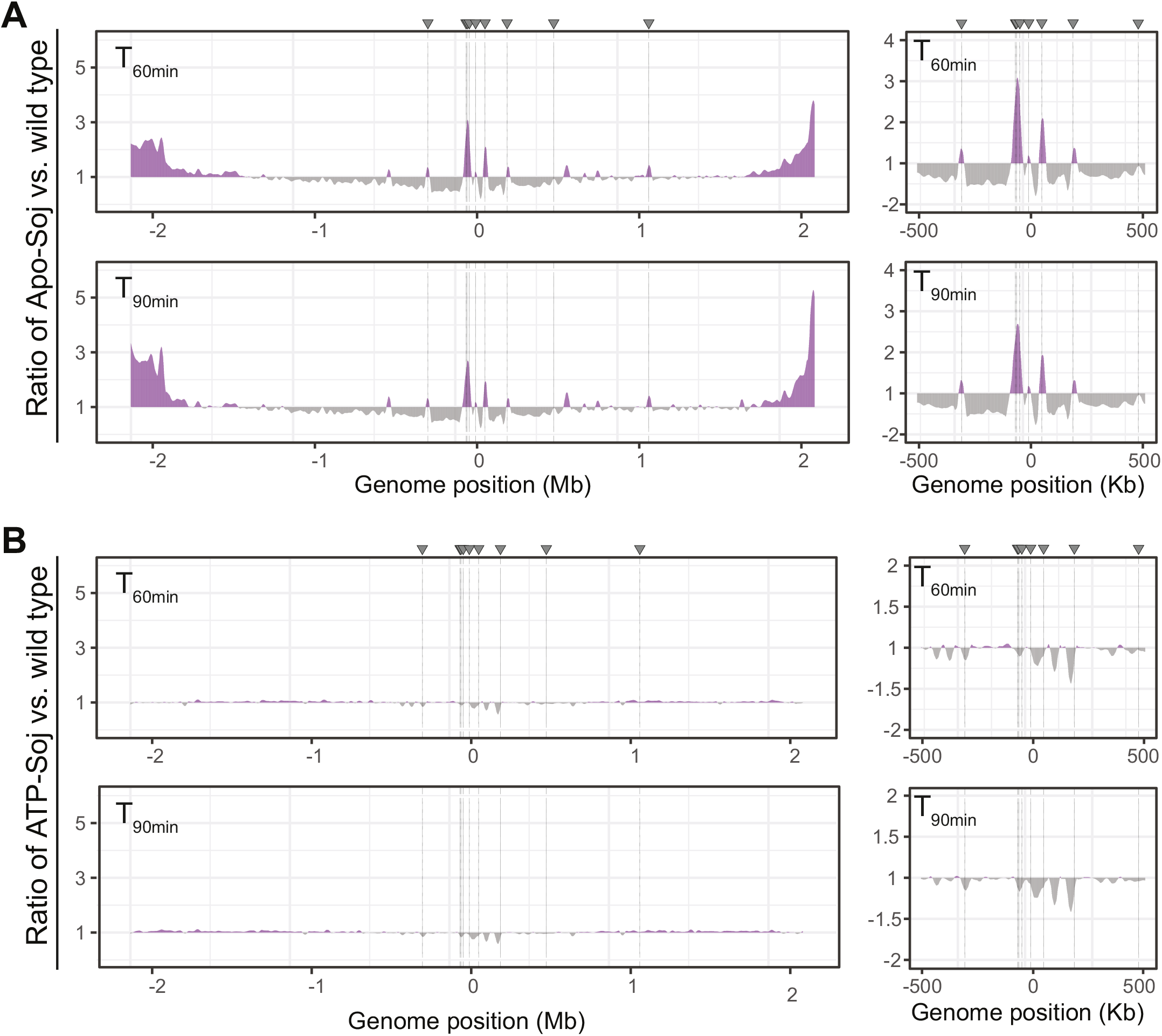
SMC complexes are differentially enriched at *parS*. ChIP-Seq was conducted against ScpB. Ratio plots show the relative enrichment (compared to wild type) of, **A)** Apo-Soj; **B)** ATP-Soj. T = time in mins after re-suspension in sporulation salts. Grey arrows and dashed lines = the 8 *oriC parS* sites. Panels to the right show a close up of the *oriC* region (+/- 500 Kb).

Supplementary Figure 6 shows the whole genome plots for each strain and condition. Interestingly, there were several peaks (green arrows) that appeared specifically after 60 and 90 minutes of sporulation, irrespective of whether the *soj* monomers were induced, suggesting that these occur when the chromosome re-organises into the axial filament during sporulation. Based on the observation of specific peaks during sporulation, we analysed the genomic context of all SMC enrichment peaks that were >4x during sporulation (t_60_ and t_90_) as compared to t_0_ (Table 1). The strongest enrichments were consistent in all conditions, and mainly corresponded to genes whose products are typically membrane integrated/associated, and that are highly transcribed in the first few hours of sporulation. In principle, these could represent roadblocks for loop extrusion by SMC complexes, similar to the foci observed by microscopy (Figure 5).

**Table 1.**
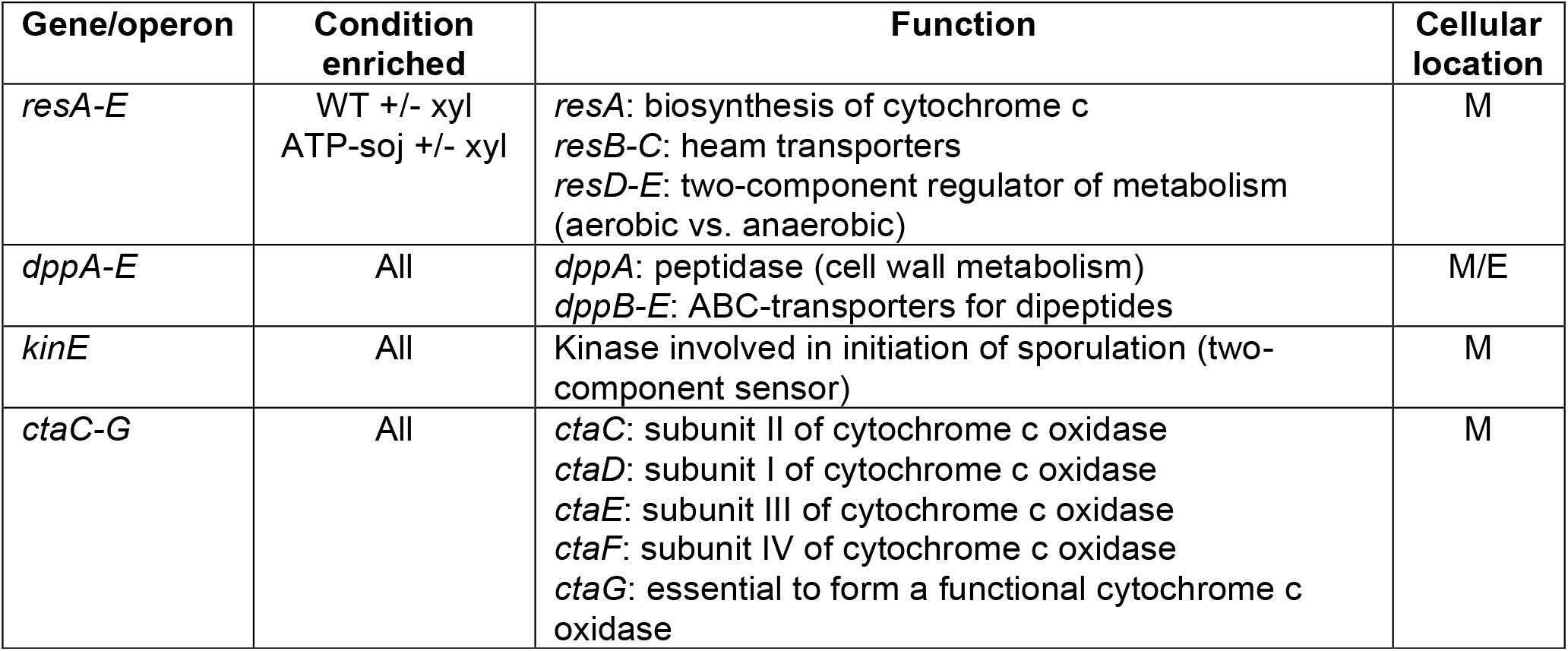
SMC is enriched along the genome during sporulation compared to vegetative growth. Table shows genes/operons at which SMC complexes were enriched >4 fold during sporulation as compared to vegetative growth (T_0 min_) based on ChIP-Seq against ScpB. M = membrane localized; E = extracellular/secreted. All information on gene function was taken and adapted from Subtiwiki (http://subtiwiki.uni-goettingen.de/)

Notably, the induced Apo-Soj ChIP-Seq profiles (Supplementary Figure 6C, +xylose, T_60_ and T_90_ mins) show no clear gradient of SMC enrichment along the chromosome arms (outside the *oriC* peaks), suggesting that there is generally less SMC translocation along the chromosome in the presence of Apo-Soj.

Upon analysing the distribution of SMC in either Apo-Soj or ATP-Soj as a ratio compared to the wild type, several striking features emerged. Firstly, there was a strong enrichment of SMC around *parS* sites in the Apo-Soj mutant (Figure 6A). Despite this increase, there was generally less SMC on the chromosome arms in the first ∼1 Mb moving away from the *oriC*. Secondly, it appeared that there was generally less SMC around the *parS* sites in the ATP-Soj mutant, but a slight increase in overall SMC localisation along the chromosome arms at both time points during axial filament formation (Figure 6B).

Taken together, these data suggest that SMC complexes are recruited to *parS* sites in the Apo-Soj mutant, but efficient translocation away (i.e. release) from the *oriC* region is blocked, ultimately leading to a depletion of SMC along the chromosome arms and a loss of the normal SMC gradients from *oriC* to *ter*. This further supports the notion that a loss of SMC along the chromosome arms is associated with an inhibition of origin segregation to the pole, which fails in Apo-Soj mutants. This is not the case in the ATP-Soj mutant, where SMC complexes are depleted around *parS* but more enriched on chromosome arms relative to the wild type, suggesting that translocation away from *oriC* is permitted and may be promoted/increased in this mutant, leading to segregation of *oriC* regions towards the cell poles.

## Discussion

### Origin segregation in sporulating *B. subtilis* does not require the dimeric form of Soj/ParA

The *soj-spo0J* locus of *B. subtilis* has long been recognized to play a role in chromosome segregation (Ireton et al., 1994; Sharpe and Errington, 1996) but its detailed functioning has been difficult to unravel, in part because of the complex phenotypic effects of mutations on chromosome replication and sporulation. Most bacteria are now known to have a *parABS* (*soj-spo0J*) system, almost always located close to *oriC* (Livny et al., 2007). The system of *C. crescentus* is probably the best understood mechanistically, as described in the Introduction. Our results clearly show that the *B. subtilis* system deviates from the *C. crescentus* paradigm, at least during sporulation. In particular, *B. subtilis* Soj/ParA appears to reside in a mainly monomeric form (Figures 1 and 2) (Murray and Errington, 2008). Moreover, the chromosome segregation defect is relatively mild, and mainly evident during sporulation when the partially redundant RacA system is abolished (Figure 3) (Kloosterman et al., 2016; Wu and Errington, 2003). Nevertheless, Soj and Spo0J appear to have all the biochemical properties needed for plasmid/*Caulobacter*-like segregation function, including the ATP-dependent dimerization and DNA binding properties of Soj. Furthermore, the *soj-spo0J* locus is sufficient to stabilize an otherwise unstable plasmid in *E. coli* (Hester and Lutkenhaus, 2007; Yamaichi and Niki, 2000). However, our results have now shown that the prominent extreme movement of *B. subtilis* origins towards the cell poles during sporulation does not require dimeric, DNA bound Soj. Why then is the dimerization and non-specific DNA binding retained? As well as a regulated switch in controlling DNA replication initiation (Murray and Errington, 2008; Scholefield et al., 2011), earlier work has shown that cells with *soj* and/or *spo0J* mutations are affected in an early step in origin separation in vegetative cells, which might account for the synthetic lethal effect observed when mutations in either gene were combined with an *smc* deletion (Lee and Grossman, 2006). Thus, it is possible that a *Caulobacter*-like DNA relay system operates to drive the initial separation of origins immediately after the initiation of replication in vegetative cells. Nevertheless, our new results suggest that the sporulation system is functionally quite different from that of the plasmid and *Caulobacter* systems.

### Apo- and ATP-bound monomers of Soj have differentiated functions

Virtually all previous work on Soj/ParA proteins has focussed on their activity in the dimeric, DNA bound state (Baxter and Funnell, 2014). However, we previously reported that an ATP-bound but monomeric form of Soj (i.e. ATP-Soj) has potent inhibitory activity against DnaA (Murray and Errington, 2008). This activity is now well established as a direct protein-protein interaction by both biochemical and structural studies (Scholefield et al., 2012; Scholefield et al., 2011). The Apo-Soj mutant has a much weaker, barely detectable inhibitory activity against DnaA (Scholefield et al., 2011), which suggested that this form of Soj might be inactive. However, our new results categorically establish that Apo-Soj is an active form of the protein: when expressed alongside the wild type it strongly inhibited sporulation, as well as generating a clear delay in nucleoid separation in vegetative cells. ATP-Soj does not have this activity. It therefore appears that neither ATP-Soj or Apo-Soj are inactive forms and, in fact, they have clearly differentiated and potent activities. Structural studies of various Soj/ParA proteins have revealed two distinct conformational states (Leonard et al., 2005; Zhang and Schumacher, 2017) and it seems possible that the ATP-Soj and Apo-Soj proteins may mimic these alternative states. Furthermore, our data suggest that the cellular functioning of wild type Soj will involve regulated switching between these states.

### Involvement of SMC/Condensin in axial filament formation and its regulation by Soj/ParA

Formation of the axial filament has long been recognized as the first detectable morphological change following the onset of endospore formation in *Bacillus* (Bylund et al., 1993; Kay and Warren, 1968; Ryter, 1965), but the mechanisms underlying this process have proven difficult to dissect. A key early finding was that *oriC* regions of the chromosome migrate towards the extreme cell poles as the axial filament forms (Glaser et al., 1997). This led to models in which the migration of origins towards opposite cell poles was responsible for the elongation of the chromosomes (Errington et al., 2005). Several factors involved in migration of the *oriC* region to the pole have been identified, particularly RacA (Ben-Yehuda et al., 2003; Wu and Errington, 2003) and a series of proteins that appear to work via an interaction with the *soj-spo0J-parS* system (Duan et al., 2016; Kloosterman et al., 2016; Wu and Errington, 2003). Until now we had assumed that the latter system would work by a kind of DNA relay mechanism involving dimeric DNA bound Soj and the ATPase stimulating activity of Spo0J. However, our new results show that Soj does not need to dimerize to support origin movement, and we have identified a change in the distribution of SMC, regulated at least in part by Soj, as the likely driving force for axial filament formation and origin migration towards the cell poles.

The change in distribution of SMC during sporulation is evident as an increase in the number of foci, which become distributed along the length of the cell (Figure 5) (Wang et al., 2015). It is well established that in the axial filament there are two complete chromosomes, with origin regions at each pole and the termini maintained close together at about mid-cell (Bogush et al., 2007; Bylund et al., 1993; Glaser et al., 1997; Hauser and Errington, 1995; Willis et al., 2020; Wu and Errington, 2003). Origin-associated SMC foci are thought to occur because this is where they are loaded (Gruber and Errington, 2009; Sullivan et al., 2009). SMC complexes then migrate away towards the terminus, meanwhile aligning the left and right chromosome arms (Wang et al., 2017; Wang et al., 2018; Wang et al., 2015). The intermediate foci seen in the axial filament might represent arm localized “pause sites”, which correlate with sporulation specific peaks of SMC enrichment observed by ChIP-Seq. These peaks occurred at the locations of genes encoding transmembrane or secreted proteins and protein complexes that are highly transcribed during sporulation. Coupled transcription-translation and membrane insertion could conceivably generate roadblocks that arrest or delay SMC loop extrusion. It will be interesting to explore whether these accumulations contribute to the changes in chromosomal topology associated with axial filament formation.

Importantly, failure of SMC to achieve this sporulation-specific chromosomal redistribution upon Apo-Soj expression was associated with a block in polar septation, suggesting that chromosome dynamics contribute to the switch in positioning of the division septum during sporulation. SMC foci have also been observed along the length of fully replicated *C. crescentus* chromosomes, which also radiate across the cell and have origins anchored at opposite cell poles (Jensen and Shapiro, 2003). It may be that distribution of SMC along the chromosome is crucial for forming an elongated nucleoid structure that traverses the entire cell axis.

The model proposed in Figure 7 illustrates one way to reconcile the possible roles of Apo-Soj and ATP-Soj in regulating SMC redistribution. It is well established that SMC complexes are loaded onto the chromosome at the origin region by Spo0J bound around *parS* sites in many bacterial species (Böhm et al., 2020; Gruber and Errington, 2009; Jalal and Le, 2020; Minnen et al., 2011; Sullivan et al., 2009). It has recently been proposed in that loading results from a direct interaction between the SMC joint and the N-terminal CTP-binding domain of Spo0J (Bock et al., 2021). At some point, SMC complexes will become released from the loading site and translocate along the chromosome to facilitate chromosome segregation and organization, as previously mentioned (Wang et al., 2017; Wang et al., 2015). Our data suggests that Apo-Soj plays a role in regulating this process, where it prevents the release of SMC complexes from Spo0J-*parS* (Figure 7, red arrow). In turn, this leads to an increase in the amount of SMC trapped around Spo0J-*parS* (relative to wild type) (Figure 6A), and a concomitant loss of SMC redistribution along the axial filament (Figure 5), as well as a failure to segregate the chromosome to the cell pole during sporulation (Figure 3B). It remains unclear whether ATP-Soj promotes SMC release from around Spo0J-*parS* or plays no active role as a positive or negative regulator. This is because, although there is a slight reduction in the amount of SMC around *parS* sites compared to wild type (Figure 6B), we cannot discount that this is due to the experimental conditions leading to a relative lack of the Apo-Soj form (i.e. the inhibitor of SMC release) in these cells.

**Figure 7.**
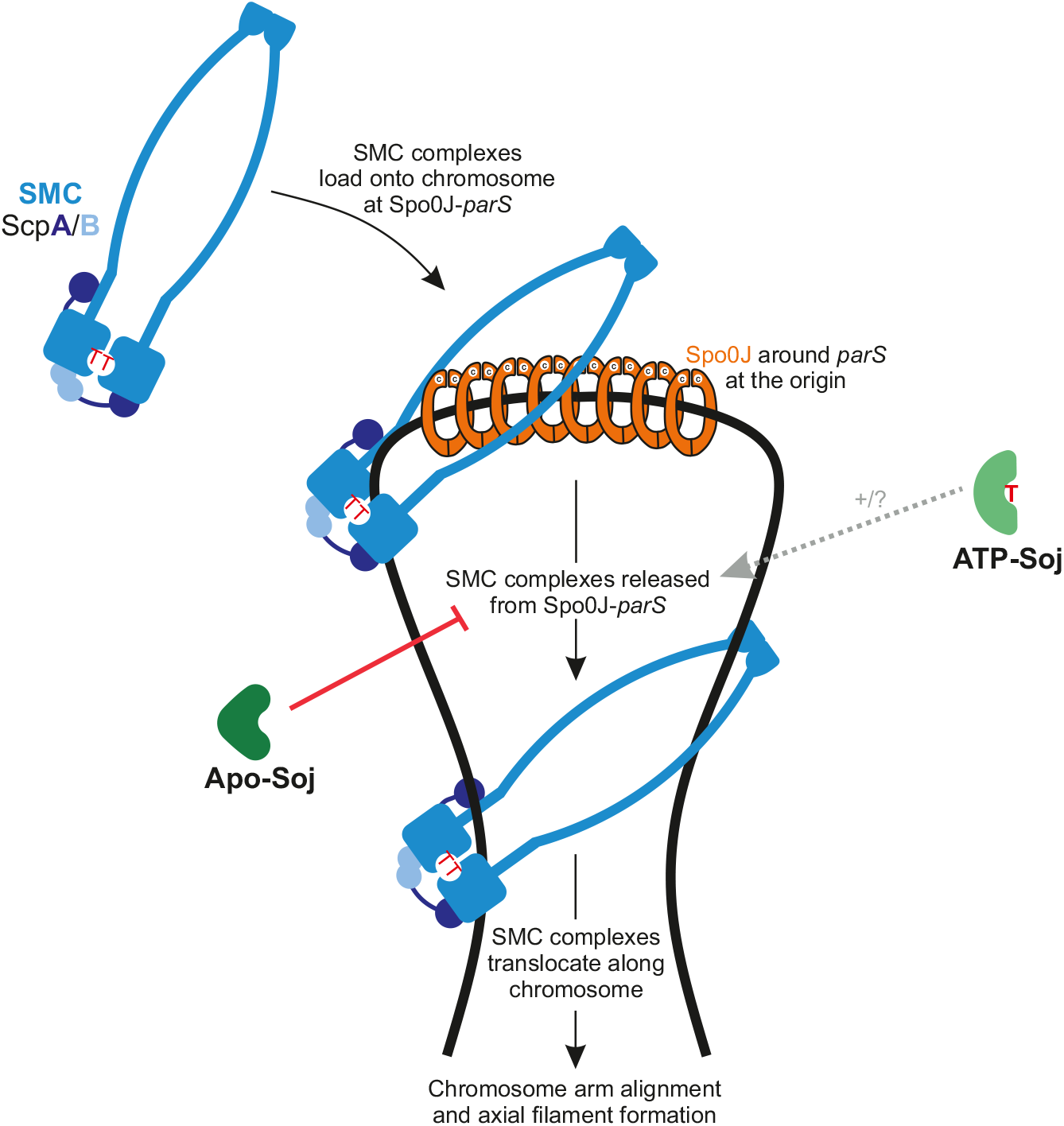
Model for the functions of Apo-Soj and ATP-Soj in controlling SMC complex redistribution in *B. subtilis*. SMC complexes are loaded onto chromosomal DNA (thick black line) by Spo0J (orange), which forms a nucleoprotein complex around *parS* sites. Once loaded, SMC complexes are released from their loading site and slide along the chromosome towards the terminus. In turn, this drives chromosome arm alignment and axial filament formation (during sporulation). Apo-Soj prevents the release of SMC complexes from Spo0J-*parS* (indicated by red line). ATP-Soj may promote the release of SMC complexes, or plays no significant role in this process (indicated by grey dashed arrow).

More work will be needed to test precisely how Soj protein regulates SMC loading and/or release into sliding mode. Nevertheless, our results appear to have identified yet another link in the complex web of interactions involving Soj, Spo0J and SMC/Condensin, which connect chromosome replication, segregation and sporulation in *B. subtilis*. They also identify new roles for both Apo- and ATP-bound forms of monomeric Soj, and provide new mechanistic insights into axial filament formation and spore septum positioning.

## Acknowledgements

We thank Maki Kawai, James Grimshaw, Alan Koh, Stijn Peters and Charles Winterhalter for strains, plasmids, and oligonucleotides; Henrik Strahl for discussions and advice on imaging and analysis; and Lizah van-der-Aart for graphic design. This work was supported by a Wellcome Investigator Award (209500 to J.E.). and the Swiss National Science Foundation (197770 to S.G.). Research support was provided to H.M. by a Wellcome Trust Senior Research Fellowship (204985/Z/16/Z).

D.M.R. was supported, in part, by the Barbour Foundation.

## Author Contributions

All authors designed the research. H.M. constructed strains. T.G.K. constructed strains and conducted initial experiments on the project. A.A. carried out the immunoprecipitations and analysis. D.M.R. conducted all other experiments and constructed the Figures. D.M.R., A.A., L.J.W., S.G., and J.E. analysed the data.

D.M.R. and J.E. wrote the manuscript with contributions from all authors.

## Methods

### Bacterial Strains and Growth Conditions

All experiments in this study used the model Gram positive bacterium *Bacillus subtilis*, with the genetic backgrounds 168CA and 168ED.

In general, strains were grown on nutrient agar (NA, Oxoid) for growth on plates, or in Luria-Bertani (LB) or casein hydrolysate (CH) liquid medium. *B. subtilis* strains were transformed by inducing competency as described by (Hamoen et al., 2002). Supplements included (when required): 0.1-1 % xylose,1 mM IPTG, 20 μg/ml tryptophan. Antibiotics were added to solid and liquid medium, when required, at the following final concentrations: 0.5-1 µg/ml erythromycin, 12.5-25 µg/ml lincomycin, 100 µg/ml ampicillin, 5 µg/ml chloramphenicol, 50 µg/ml spectinomycin, 2-5 µg/ml kanamycin, 6-10 µg/ml tetracycline, 10 µg/ml zeocin.

### *Sporulation of* B. subtilis

Strains of *B. subtilis* were induced to sporulate using the resuspension method (Harwood and Cutting, 1990; Partridge and Errington, 1993; Sterlini and Mandelstam, 1969). In summary, cultures (3-10 ml) were inoculated from fresh plates and grown overnight at 30°C in CH medium containing the appropriate antibiotic. The following day, cultures were diluted to an optical density (OD_600_) of 0.1 in fresh CH medium and grown at 30°C or 37°C until the OD_600_ reached 0.8. Cultures were then pelleted and resuspended into an equal volume of A+B sporulation salts, which marked t_0_ of sporulation. All subsequent time points for sporulation (in min or h) are measured similarly (e.g. t_90 min_ marks 90 minutes after re-suspension in sporulation salts).

### Microscopy

In all microscopy experiments, cells were imaged using a Nikon Ti microscope equipped with a Nikon CFI Plan Apo DM Lambda x100 oil objective; MetaMorph v7.7 software (Molecular Devices); Photometrics Prime or BSI SCMOS cameras; and a Sutter Instruments Lamda LS xenon arc or CoolLED PE-300/PE-4000 LED light sources. Images were processed and analysed using FIJI (https://imagej.net/Fiji) (Schindelin et al., 2012).

### Single time point fluorescence microscopy from batch cultures

For single time point imaging, overnight cultures of vegetatively growing cells were diluted to OD_600_=0.05-0.1 and grown for at least two cell doublings prior to imaging. For sporulation experiments, cells were imaged at appropriate times following re-suspension into sporulation salts (see Figures for details). In all cases, 0.5 µl cells were mounted onto microscope slides containing a thin layer of 1 % agarose in water (supplemented with 10 % medium, where appropriate) and covered with 0.13-0.17 mm coverslips (VWR). Nucleoids were visualised using 1 µg/ml DAPI (Sigma Aldrich) and membranes were visualised by adding 0.5 µg/ml FM5-95 (Molecular Probes) to the agarose pad or cells. To minimise the non-specific binding of FM5-95, coverslips were pre-coated with a large droplet of polydopamine (Sigma Aldrich) (2 mg/ml dopamine in 5 mM Tris-HCl, pH 8.5) prior to sequential washing (twice) in deionised water (Te Winkel et al., 2016).

### Chromosome Trapping Assay during Sporulation

Three hours after the induction of sporulation by the resuspension method, cells were analysed by microscopy. Prespores, identified by membrane staining with FM5-95, were scored for the expression of YFP and/or CFP and categorised as expressing both markers, YFP only, CFP only, neither marker, or as other (denoting any other pattern e.g. expression of markers in the wrong/both compartments). The presence of a specific point mutation in SpoIIIE (*spoIIIE36*) prevented the translocation of the bisected prespore chromosome from its initial capture orientation, allowing the expression and maturation of the prespore-specific fluorescent markers.

### Image Analysis

In all cases, FIJI software (https://imagej.net/Fiji) (NIH) was used to process and analyse imaging data (Schindelin et al., 2012).

### SMC-mNG spot detection

Individual SMC-mNG foci were assigned as “spots” in a semi-automated, non-biased manner by using the TrackMate plugin in FIJI (Tinevez et al., 2017). In TrackMate, the Laplacian of Gaussian filter was applied, and an estimated blob diameter of 5 pixels and a threshold of 250 grey levels was used to assign all SMC-mNG spots in a field of view. Following background subtraction and generation of a composite image merging the green and red (i.e., SMC-mNG and membrane) channels, the assignment of individual spots on the merge image was generated. The channel representing the SMC-mNG foci was selected as the active channel for spot detection in the software. The generated image was then saved, and the detected spots/cell counted manually.

### Fluorescence Profile generation for mNG-Soj and DivIVA-mScl-I

Images of mNG-Soj (WT, Apo- and ATP variants) and DivIVA-mScl-I were loaded into FIJI. Foci of Soj and DivIVA were detected as spots using the TrackMate plugin with the Laplacian of Gaussian filter. An estimated blob diameter of 7.5 and 8.0 pixels was used for Soj and DivIVA images, respectively, and a background threshold of 750 grey levels was applied to all images. Spots were then exported as ImageJ ROIs, which generated an image containing just the assigned spots. A mask was applied prior to image smoothing. This process was then repeated for every image, position and both Soj and DivIVA channels for every strain and time point. This removed all non-specific signal apart from the software-assigned spots that represent the position of Soj/DivIVA foci.

To generate fluorescence profiles, all phase images and the assigned Soj and DivIVA spots for each imaging field were merged into an hyperstack for each strain, generating a stack each for wild type, ATP-Soj and Apo-Soj. The images were analysed, and profiles automatically assigned to each cell using the Coli-Inspector tool (version 04d) in the ObjectJ plugin for Fiji (Vischer et al., 2015). In brief, each hyperstack of images were individually selected as the linked image in the ObjectJ Coli-Inspector tool, and then filaments were automatically marked using the software. Following the generation of sorted and qualified maps, the corresponding profiles were saved and processed manually as greyscale or pseudocoloured images.

### Chromatin Immunoprecipitation coupled with deep sequencing (ChIP-Seq)

For the single time point ChIP-Seq, 10 ml overnight cultures of strains DMR178, DMR179 and DMR181 grown in CH media were diluted to OD_600_ = 0.1 in 50 ml fresh CH. Each strain was then grown and induced to sporulate by the resuspension method (described above). After 3 hours, each sample was crosslinked with formaldehyde (1 % final) at room temperature for 30 min with shaking every 10 mins. Crosslinked pellets were harvested by centrifugation and washed twice in PBS prior to rapid freezing in liquid nitrogen and storage at −80°C until further processing For the time-course ChIP-Seq, 10 ml overnight cultures of strains DMR312, DMR314 and DMR363 grown in CH media were diluted to OD_600_ = 0.1 in 100 ml fresh CH. Strains were then induced to sporulate by the re-suspension method (as above).

Upon re-suspension in A+B sporulation salts, each sample was split into 2x 50 ml samples, with expression of the ectopic copy of *soj* being induced in one of each pair by the addition of 0.5 % (final) xylose. Each sample was handled independently throughout. At t_0 min_, t_60 min_ and t_90 min_ of sporulation, 15 ml was removed from each culture and crosslinked with formaldehyde (1 % final) for 30 min at room temperature with shaking every 10 mins. Each sample pellet was washed twice in PBS, prior to rapid freezing and storage at −80°C until further processing.

Each sample pellet was resuspended in 2 ml cold PBS and adjusted to 4 OD_600_ units (4 ml at OD_600_=1). Adjusted sample pellets were resuspended in TSEMS buffer (50 mM Tris pH 7.4, 50 mM NaCl, 10 mM EDTA pH 8.0, 0.5 M sucrose and PIC (Sigma)) supplemented with 10 mg/ml lysozyme from chicken egg white (Sigma) and incubated for 30 min at 37°C with vigorous shaking. Resulting protoplasts were collected by centrifugation, washed twice with TSEMS, resuspended in 1 ml TSEMS and split into 3 aliquots. Pelleted samples were rapidly frozen in liquid nitrogen for storage at −80°C until further processing.

For immunoprecipitation (IP), the sample pellets were resuspended in 2 ml buffer L (50 mM HEPES-KOH pH 7.5, 140 mM NaCl, 1 mM EDTA pH 8.0, 1 % (v/v) Triton X-100, 0.1 % (w/v) Na-deoxycholate, 0.1 mg/ml RNaseA and PIC (Sigma)) and transferred to 50 ml round bottom tubes. Sonication was performed using a Bandelin Sonoplus with an MS72 tip (90 % pulse and 35 % power output) for 3 rounds of 20 sec. Lysates were transferred to 2 ml tubes and, after centrifugation for 10 min at 21,000 g at 4°C, 800 μl of the supernatant was collected for IP and 200 μl was kept as whole-cell extract (WCE) at −20°C.

Prior to IP, the anti-scpB antibody serum (Gruber Lab) was incubated with Protein G coupled Dynabeads (Invitrogen) in a 1:1 ratio for 2.5-3.5 hrs at 4°C with rotation. The antibody-bead mixture was then washed and resuspended in buffer L, prior to adding 50 μl to each IP sample, which were incubated for 2.5-3 hrs at 4°C with rotation. Next, the samples were washed with the following buffers: buffer L, buffer L5 (buffer L containing 500 mM NaCl), buffer W (10 mM Tris-HCl pH 8.0, 250 mM LiCl, 0.5 % (v/v) NP-40, 0.5 % (w/v) Na-deoxycholate, 1 mM EDTA pH 8.0), and buffer TE (10 mM Tris-HCl pH 8.0, 1 mM EDTA pH 8.0). Finally, for crosslink reversal, the IP samples were resuspended in 520 μl buffer TES (50 mM Tris-HCl pH 8.0, 10 mM EDTA pH 8.0, 1 % (w/v) SDS) and transferred to 1.5 ml screw-cap tubes. WCE samples were thawed and 300 μl of TES and 20 μl of 10 % SDS were added. Both tubes were incubated overnight at 65°C with vigorous shaking.

To purify the DNA, 500 μl phenol:chloroform:isoamyl alcohol mix (Sigma) was added to the IP and WCE tubes, mixed vigorously and centrifuged for 10 min at 13,000 rpm (at room temperature). 450 μl of aqueous phase was collected and precipitated with 1 ml 100 % ethanol in the presence of 45 μl NaOAc (Sigma) and 1.2 μl of Glycoblue (Invitrogen) for 20 min at −20°C. DNA pellets were collected by centrifugation and resuspended in 100 μl EB (Qiagen) by vigorous shaking for 10 min at 55°C. Final purification was performed using a PCR purification kit (Qiagen), eluting the DNA in 50 μl EB. Prior to deep sequencing, the success of the IP was verified with qPCR.

For deep sequencing, the DNA libraries were prepared by the Genomic Facility at CIG, UNIL, Lausanne. Briefly, the DNA was fragmented by sonication (Covaris S2) until the DNA was sheared to 220-250 bp. The Ovation Ultralow Library Systems V2 kit (NuGEN) including 15 cycles of PCR amplification was used to prepare the DNA libraries. 12-15 million sequence reads per sample were obtained on a HiSeq4000 (Illumina) with 150 bp read length.

### ChIP-Seq read profiling

The *B. subtilis* genome NC_000964.3 was used to map the reads using bowtie2 (--very sensitive-local mode). Downstream data analysis was conducted using SeqMonk (Babraham Institute), with a bin size of 1 kb. Data was visualized using R or GraphPad Prism.

