## Supplemental Data for "Chromosome remodelling by SMC/Condensin in *B. subtilis* is regulated by Soj/ParA during growth and sporulation"

### Supplementary Figure 1

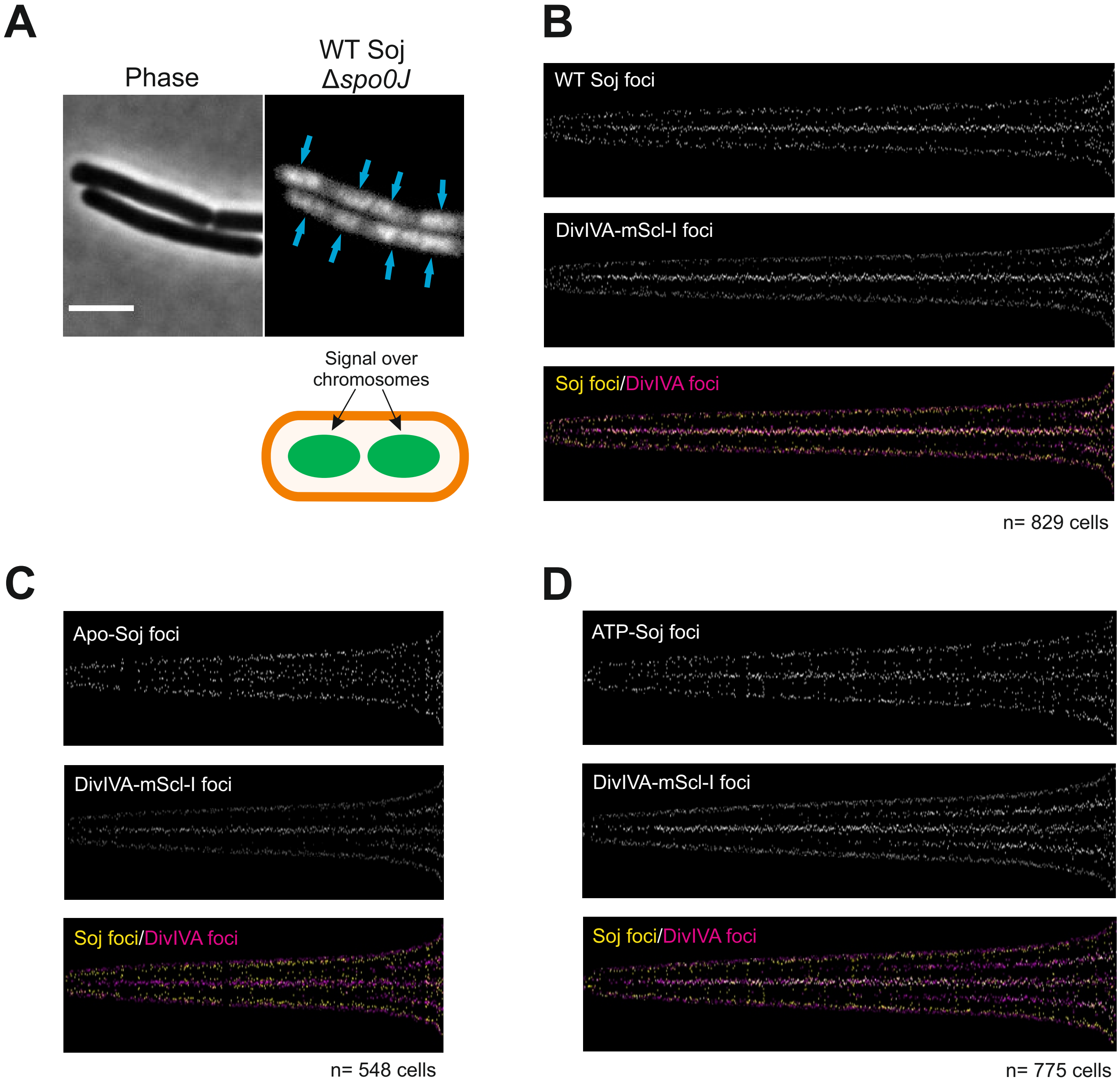


**Supplementary Figure 1. Analysis of Soj and DivIVA localization**
**A)** Representative image showing the localization of mNG-Soj in a Δ*spo0J* mutant. Blue arrows = localization of Soj over the chromosomes. Scale bar = 3 µm. **B-D)** Fluorescent plots showing the relative cellular positions of **B)** wild type mNG-Soj and DivIVA-mScl-I; **C)** Apo-Soj and DivIVA-mScl-I; **D)** ATP-Soj and DivIVA-mScl-I. In all cases, cells were sorted by shortest to longest. The DivIVA-mScl-I signal was used as a proxy for the cell boundaries/poles.

### Supplementary Figure 2

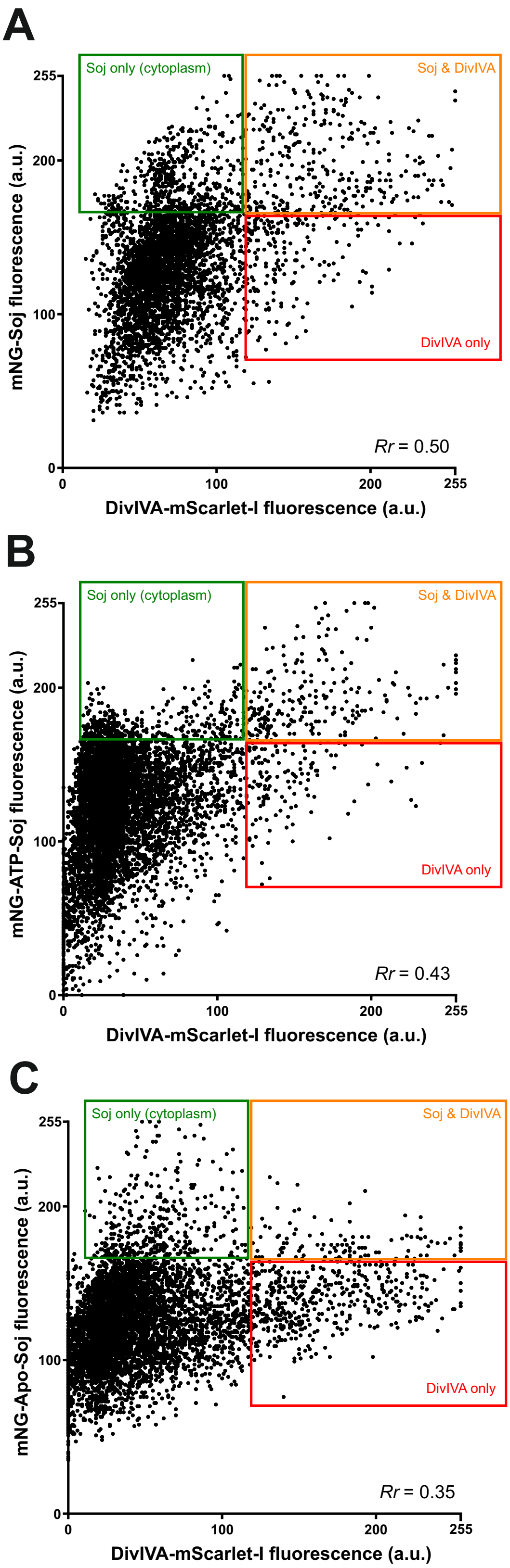


**Supplementary Figure 2. Co-localization analysis of mNG-Soj and DivIVA-mScl-I**
The graphs show a fluorescence intensity correlation (pixel by pixel) of the Soj and DivIVA signal for **A)** wild type Soj and DivIVA; **B)** ATP-Soj and DivIVA; **C)** Apo-Soj and DivIVA. Images used are those from Figure 2. The Pearson's correlation coefficient (*Rr*) is also shown. Green boxes = pixels corresponding to Soj only signal; orange boxes = pixels corresponding to co-localized Soj and DivIVA; red boxes = pixels corresponding to DivIVA only signal.

### Supplementary Figure 3

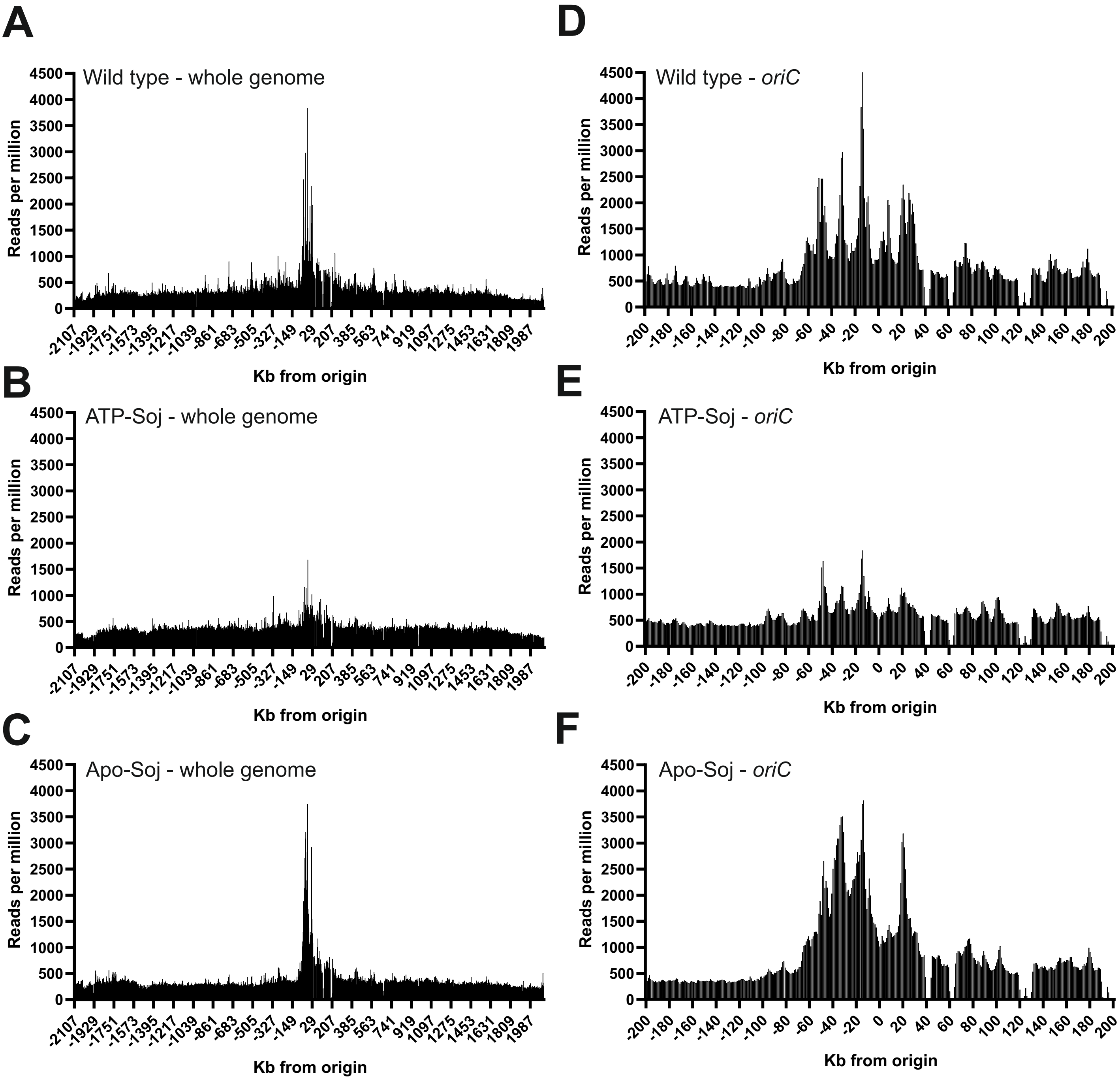


**Supplementary Figure 3. ChIP-Seq reveals altered enrichment of SMC at *oriC* during sporulation**

ChIP-Seq against ScpB was conducted 3 h after re-suspension in sporulation salts. **A-C)** Whole genome ChIP-Seq profiles are shown for **A)** wild type; **B)** ATP-Soj; **C)** Apo-Soj. **D-F)** a close up of the origin region (+/- 200 Kb) is shown for **D)** wild type; **E)** ATP-Soj; **F)** Apo-Soj. To lock the chromosome in its initial capture orientation, all strains also contained the *spoIIIE36* mutation.

### Supplementary Figure 4

**A**

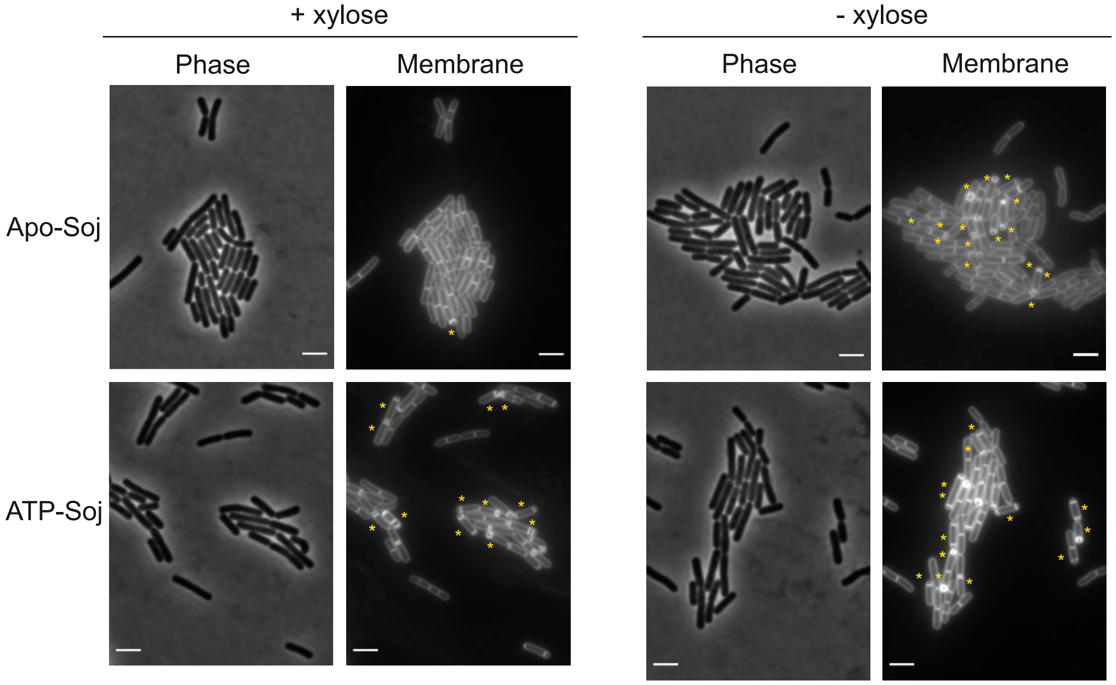


**B**

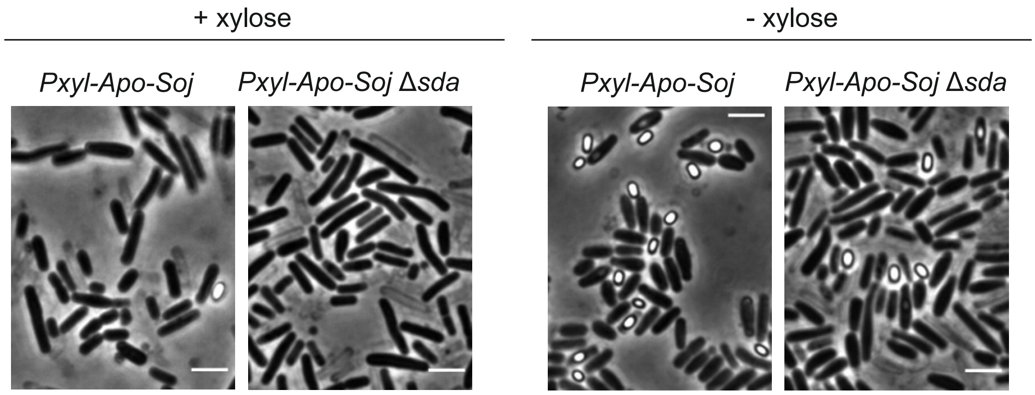


**C**

| Time Point during sporulation | Asymmetric Division | Mean intensity of $P_{spoIIA}$-*mcherry* |
|---|---|---|
| ATP-Soj +xyl, 240min | Yes | <u>684</u> |
| ATP-Soj –xyl, 240min | Yes | 2018 |
| Apo-Soj +xyl, 240min | No | <u>618</u> |
| Apo-Soj –xyl, 240min | Yes | 2005 |

**Supplementary Figure 4. The Apo-Soj monomer is dominant negative over wild type.**
**A)** Representative images showing cells in which Apo-Soj (top panels) and ATP-Soj (bottom panels) were either overexpressed (+ xylose) or not (- xylose) 100 mins after re-suspension in sporulation salts. For + xylose, 0.5 % xylose was added 15 mins prior to (and during) re-suspension in sporulation salts. Membranes were visualized using FM5-95. Yellow asterisks show examples of cells containing asymmetric septa. Scale bar = 3 µm. **B)** Representative images of cells showing the dominant negative effect is not lost in Δ*sda.* + xylose plates contained 0.5 % xylose. - xylose = no induction of Apo-Soj. Scale bar = 3 µm. **C)** Representative fluorescence intensities showing the induction of a PspoIIA-mcherry marker. + xyl = addition of 0.5 % xylose to cultures 15 mins prior to (and during) re-suspension in sporulation salts. -xyl = no induction of ATP- or Apo-Soj.

### Supplementary Figure 5

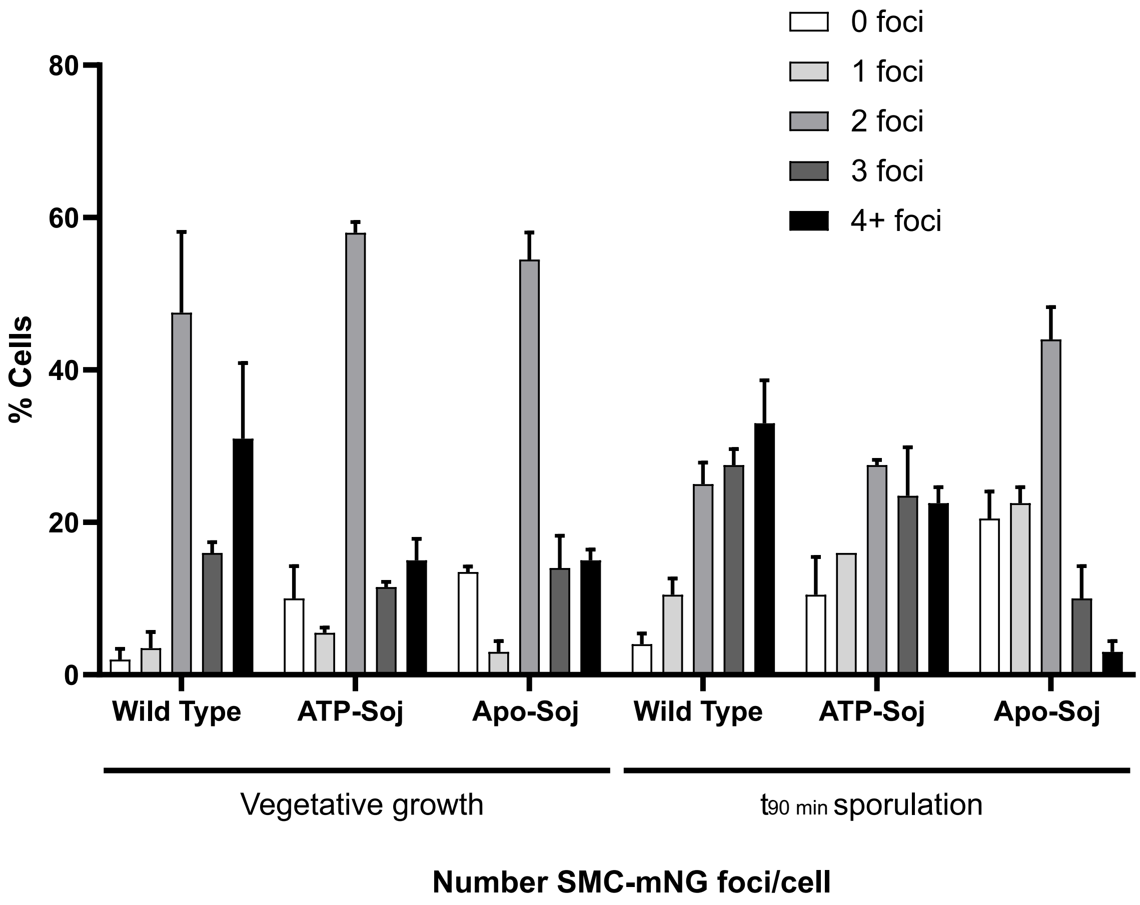


**Supplementary Figure 5. SMC complex redistribution is altered during sporulation.** Bar chart showing the number of SMC-mNG foci per cell in wild type, ATP-Soj and Apo-soj mutants expressed as single copies from their native locus. At least 100 cells were counted at each time point (n = 2) and average values are shown. Error bars = standard deviations.

### Supplementary Figure 6

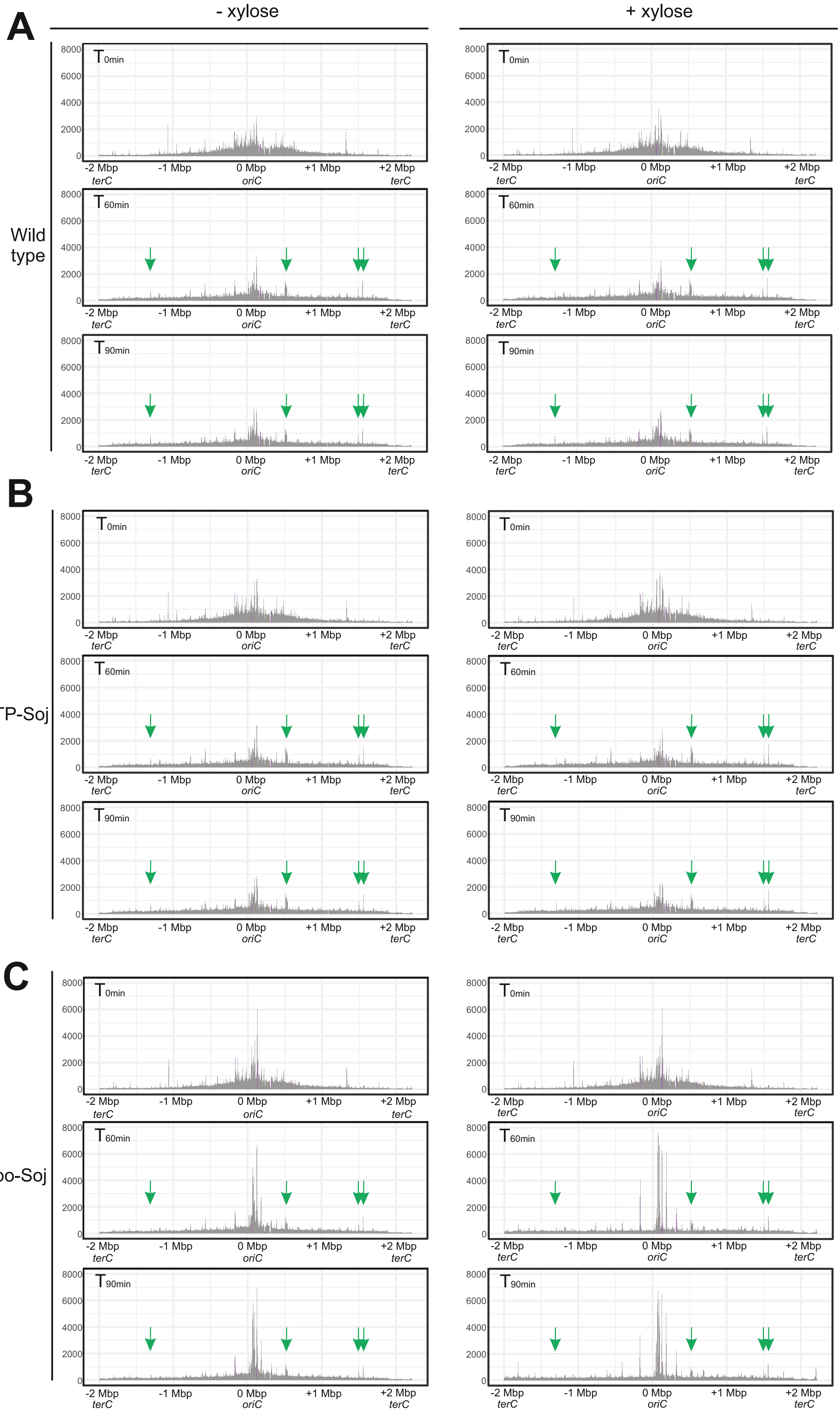

**Supplementary Figure 6. Whole genome anti-ScpB ChIP-Seq plots during early sporulation.** **A)** wild type; **B)** ATP-Soj; **C)** Apo-Soj. Cells were sporulated in the presence (+ xylose) or absence (- xylose) of 0.5 % xylose. T = minutes post re-suspension in sporulation salts. Green arrows = peaks specific to sporulation in all conditions. Purple lines = *parS* sites. Plots show reads per million.
